# Maize stigmas react differently to self-/cross-pollination and fungal invasion

**DOI:** 10.1101/2024.04.17.589941

**Authors:** Kevin Begcy, Mariana Mondragón-Palomino, Liang-Zi Zhou, Patricia-Lena Seitz, Mihaela-Luiza Márton, Thomas Dresselhaus

## Abstract

During sexual reproduction in flowering plants, rapid tip-growing pollen tubes travel from the stigma inside the maternal tissues of the pistil towards ovules. In maize, the stigma is highly elongated forming thread-like strands known as silks. Only compatible pollen tubes are supported to penetrate and to grow successfully through the transmitting tract of the silk to reach the ovules. Like pollen, fungal spores germinate at the surface of silks and generate tube-like structures (hyphae) penetrating silk tissue. To elucidate commonalities and differences between silk responses to various invaders, we compared growth behavior of the various invaders as well as the silk transcriptome after self-pollination, cross-pollination and infection using two different fungi. We report that self-pollination triggers especially senescence genes, while incompatible pollination using *Tripsacum dactyloides* leads to downregulation of rehydration genes, microtubule and modulation of cell wall-related genes explaining slower pollen tube growth and their arrest. Invasion by the ascomycete *Fusarium graminearum* triggers numerous defense responses including the activation of monolignol biosynthesis and NAC as well as WRKY transcription factor genes, while responses to the basidiomycete *Ustilago maydis* are generally much weaker. Cell wall and phytoalexin biosynthesis pathways were selected as examples to demonstrate usability of the data sets provided.

**One sentence summary:** Data sets are provided and analyzed to elucidate how maize stigmas (silks) respond to different biotic invaders including own and alien pollen tubes as well as hyphae of two different fungi.

## Introduction

Maize (*Zea mays* L.) is a monoecious plant with unisexual female and male flowers (Dresselhaus et al., 2011). While the female inflorescence, also known as ear, is originating from axillary meristems, the male inflorescence or tassel is positioned at the apex of the plant and originates from the shoot apical meristem after transition to various flower meristems (Zhou et al. 2017; Bommert and Whipple 2018). Ears and tassels initially undergo a similar developmental program but eventually develop into distinct architectures. In developing ears, the lower floret is aborted, and only the upper floret continues to develop into a functional flower forming an ovary and stigma, the latter also known as silk (Abendroth et al. 2011). Maize silks are long thread-like strands exposed out of the ear shoot and function as the main pollen receptive organ. Numerous hairs (papilla hair cells or receptive trichomes) are distributed along the length of a silk and function as major surfaces for pollen capture and germination (Zhou et al. 2017). The main function of maize silks is to allow own compatible pollen to hydrate, germinate and successfully penetrate papilla hair cell structures with the goal to carry their sperm cell cargo at long distances and high-speed growth inside the transmitting tract towards the female gametophyte or embryo sac inside the ovary (Lausser et al. 2010). However, maize silks also receive pollen from other plants species and are exposed to microorganisms including pathogens. While pollen from plant species outside the grass family are partly capable to germinate on papilla hair cells, they are usually not capable to penetrate. Pollen from other grass species like the maize relative *Tripsacum dactyloides* are capable to penetrate, but growth is not supported and arrested after about 2-3 cm inside the silk tissue (Lausser et al. 2010). Like pollen grains, spores of fungal pathogens germinate on silks generating tip-growing tubes or hyphae and use silks as a route to ultimately invade developing kernels (Thompson and Raizada 2018). Fungal hyphae growth inside silks is significantly slower compared to pollen tube growth and further inhibited if silks were simultaneously exposed to compatible pollen (Bircheneder and Dresselhaus 2016; Thompson and Raizada 2018). Flowering plants have evolved sophisticated molecular responses to reject alien (incompatible) pollen and pathogens. While similarities of recognition mechanisms between alien pollen and pathogens have been proposed to be widespread in plants, only few studies have compared both processes (Kessler et al. 2010; Mondragón-Palomino et al. 2017). Most of the common associated responses have been narrowed down to receptor-like kinases (RLKs) and phytohormones during defense mechanisms and pollen incompatibility (Mondragón-Palomino et al. 2017; Shi et al. 2017). Although our understanding of how plants (especially Arabidopsis) respond to pollination (Sella Kapu and Cosgrove 2010; Mondragón-Palomino et al. 2017; Shi et al. 2017; Mandrone et al. 2019; Rozier et al. 2020; Kodera et al. 2021) and pathogens (Mondragón-Palomino et al. 2017; Ning and Wang 2018; Zhou et al. 2018; Li et al. 2019; Yadav et al. 2020; Kumar et al. 2022) has improved significantly in recent years, it is unclear whether both processes trigger similar or different responses and comparable studies of pollination and pathogen infection in maize have to our knowledge not been performed.

We used maize as a model in this study as its elongated stigmas (silks) are easily accessible, can be harvested in large quantities and allows to follow the invasion of different biotic invaders over long distances and time. GFP-lines were used to follow the growth of pollen tubes and fungal hyphae, respectively. A dataset of 11 different samples was generated including also aging and vegetative tissue controls. To identify common, different and specific responses towards the various invaders, we compared molecular signatures of maize silks after exposure to fungal infections using the hemi-biotrophic ascomycete *Fusarium graminearum* and the biotrophic basidiomycete *Ustilago maydis* as well as compatible pollination (self-pollination) and incompatible (cross-) pollination using the close maize relative *Tripsacum dactyloides*. While expression of relative few genes was changed after self-pollination, cross-pollination and especially infection by *F. graminearum* led to drastic gene expression changes. A number of processes especially related to genes involved in cell wall and lignin biosynthesis and modification, gene regulation as well as phytoalexin production has been studied in more detail.

## Results and Discussion

### Maize silks are direct highways used by pollen tubes and fungal hyphae to enter ovaries

The elongated silks of maize represent an ideal cereal model system to explore and compare cellular and molecular strategies used by invading pollen tubes and pathogens during reproduction and defense responses, respectively. Even though some similarities between reproductive and defense responses have been reported in *Arabidopsis* (Mondragón-Palomino et al. 2017), it is unclear to which extent this can be translated to cereals and whether responses of the silk to incompatible pollen is similar to fungal pathogen invasion. To test this hypothesis, we used the fungal pathogens *F. graminearum* and *U. maydis* to study defense responses. To elucidate reproductive responses, we used pollen of the maize relative *T. dactyloides* and *Zea mays* (maize) as incompatible and compatible material, respectively (Figure 1A). First, we analyzed invasion of pollen tubes. Silks were pollinated using maize pollen carrying a GFP marker (Figure 1B-E). After adhesion, pollen grains germinated within about 10 minutes and penetrated papilla hair cell structures between cell boundaries. Further growth occurred intercellularly in the center of hair cell structures (Figure 1B, C) by pushing cells apart (Figure 1D). After entering the main body of the silk, maize pollen tubes grew intercellularly towards the transmitting tract (Figure 1E). While *T. dactyloides* pollen tubes entered papilla hair cell structures using the same strategy, they often did not enter the transmitting tract and growth was arrested after 2-3 cm growth as reported previously (see also Lausser et al., 2010).

**Figure 1.**
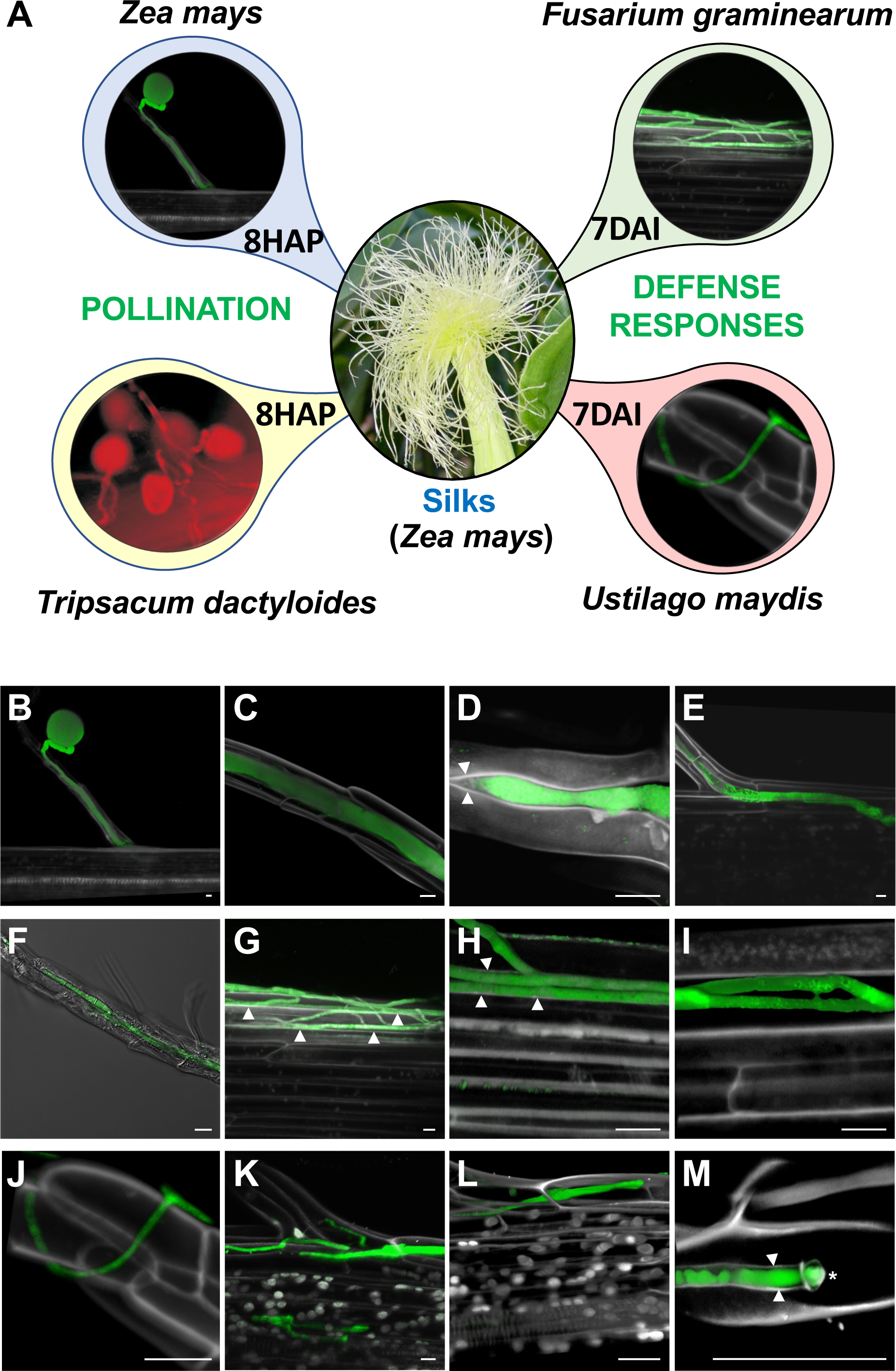
Invasion of papilla hair cells and silks of maize by own and alien pollen tubes as well as by two different fungi. **(A)** Experimental setup: maize silks were pollinated with *Zea mays* B73 (self-pollination) and *Tripsacum dactyloides* (incompatible pollination), respectively, and samples collected at 8 hours after pollination (HAP). Additionally, silks of maize plants were infected with the facultative hemibiotrophic pathogen *Fusarium graminearum* and the biotrophic pathogen *Ustilago maydis*. Samples were collected at 7 days after infection (DAI). **(B-E)** A GFP-labeled maize pollen tube germinated at the apex of a papilla hair cell structure and grows through the papilla (B) in the intercellular space between hair cells (C) while pushing hair cells apart (D) and growing straight inside the silk towards the transmitting tract (right) after reaching into the bottom of the hair cell structure. Arrowheads indicate cell walls. **(F-I)** GFP expressing *F. graminearum* hyphae invade silks either after penetration of papilla hairs and intercellular growth like pollen tubes (F) or after initial growth at the silk surface (G). Growth inside the silk occurs first intercellular by pushing cell walls apart (H) before cells are penetrated and die (I). Arrowheads indicate cell walls. **(J-M)** GFP-labelled *U. maydis* growth through the silk ‘ignoring’ cell walls. An appressorium is visible at the surface of a papilla hair (J) and further fungal growth occurs intracellularly (K-L). Cell wall material is always detectable surrounding intracellular growing fungal hyphae (M). The tip of a hyphae is marked by an asterisk. The white color in all images results from propidium iodide counterstaining. Bars: 10 μm.

The overall pattern of silk infection was investigated for both pathogens using also GFP marker lines. Macroconidia or ascospores of *F. graminearum* germinated on exposed silks and entered silk hair cell structures using the intercellular pollen tube route and by pushing cells apart. After entering the body of the silk, further hyphae growth occurred initially intercellularly, before cells were penetrated and killed. Later growth occurred intracellular (Figure 1F-I). *U. maydis* entered maize silks after forming appressoria (Figure 1J). Further growth occurred exclusively intracellularly without killing cells (Figure 1K and L). Fungal hyphae were always surrounded within cells by a plasma membrane and thin cell wall structures (Figure 1M).

### Pathogen infection and foreign pollination elicit unique and only slightly overlapping pattern of molecular responses in maize silks

To characterize the reproductive and defense responses at the molecular level, we explored the transcriptional changes in all above-described scenarios. Silks of the maize inbred line B73 were self-pollinated and analyzed 8 hours after pollination (8HAP) using own B73 pollen (referred to hereafter as “Silks self-poll”). Incompatible pollination was studied after pollinating maize silks with *T. dactyloides* pollen (“Silks *Tripsacum*”). As controls, we included mature unpollinated silks (“Silks un-poll”) of the same age (8H) and pollen tubes collected after 45 minutes (45M) *in vitro* germination (“Pollen tubes”) in order to capture silk-specific responses. To elucidate fungal defense responses, maize silks were infected either with *F. graminearum* (“Silks *Fusarium*”) or *U. maydis* (“Silks *Ustilago*”) and material was collected 7 days after infection (7DAI). Two types of controls were included: mature silks 7 days (7D) after mock-treatment (“Silks mock”) and untreated mature silks 7 days (“Silks untreated”) as aging control. As additional controls for both groups, we included seeds, leaves, and roots of maize to elucidate silk-specific responses. Supplemental Table 1 summarizes the sample material, average of total reads per condition, and percentage of mapped reads. A detailed list containing the median expression values of treatment/tissue for the entire maize genome is provided in Supplemental Data set 1.

A principal component analysis (PCA) of all samples showed clear separation of the different tissues and close association of the three biological replicates used per sample for RNA-seq analysis. This indicated that the samples were highly reproducible. PCA on the gene expression data showed that PCA1 largely separates silk samples from pollen tubes, seeds, and roots (Figure 2A). To validate RNA-seq data, we randomly selected 33 genes that were expressed in more than one treatment (Supplemental Figure S1). We found a 0.823 correlation coefficient between the RNA-seq results and the real-time PCR data confirming the high quality and reproducibility of the data. A comparison of all genes expressed in silks under the different conditions studied showed similar percentages of total transcriptional expression (about 55% of all maize genes). A comparative percentage of expression was observed in seeds, leaves, and roots. Notably, pollen tubes expressed only about 20% of all maize genes (Figure 2B). This can be explained by the fact that they consist of only two cell types of highly specific functions compared to the other very complex tissues containing many different cell types. We further identified clusters of genes characteristic for all samples (Figure 2C). While silks unpollinated and self-pollinated showed a similar pattern, the pattern after *T. dactyloides* pollination was already quite different. Infection with *F. graminearum* triggered the largest response in maize silks. Noteworthy, silks harvested 7D after mock-treatment and untreated showed a highly similar pattern, but different from young unpollinated silks indicating indeed aging effects.

**Figure 2.**
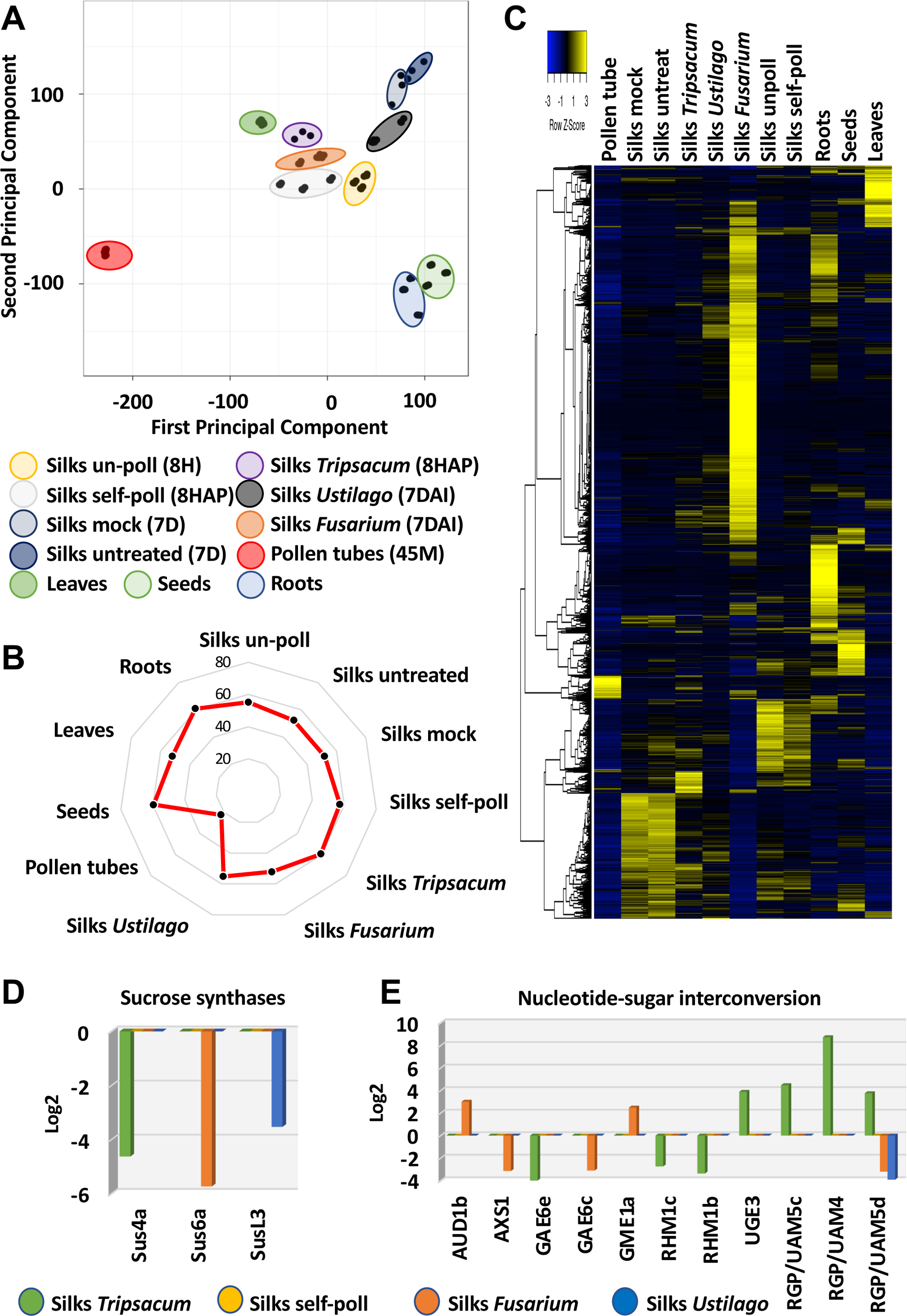
Transcriptome relationships between pollinated and infected maize silks. **(A)** Principal component analysis (PCA) of RNA-seq data from the 11 tissue samples used in this study (see Supplemental Table S1 for sample overview). Three biological replicated were used for each sample that include various pollinated, infected and control samples as indicated. **(B)** Radar (spider) chart displaying the percentage of transcripts from the maize genome expressed in each tissue. **(C)** Heat Map of the whole maize genome in response to self-pollination, incompatible pollination and fungal infection. **(D)** Genes involved in sucrose synthase and **(E)** nucleotide-sugar inter conversion are strongly regulated after invasion of foreign pollen tubes and fungal hyphae, respectively.

We next studied whether there exist gene groups that are differentially regulated in response to foreign invaders, but which are not regulated by own pollen. Three sucrose synthase genes were identified, for example, with silk-preferred expression that were downregulated in response to incompatible pollination and fungal invasion (Figure 2D). While *Sus4a* was downregulated in response to *T. dactyloides*, *Sus6a* was repressed after *F. graminearum* infection and *SusL3* in response to *U. maydis*. Sucrose synthases are considered as markers for sink strength in crops (Xu et al. 2019), thus their down-regulation may indicate a reduction of starch, cellulose or callose biosynthesis, but also reduced tolerance to stress conditions (Stein and Granot 2019) and biotic invaders.

One of the first steps during cell wall biosynthesis is substrate generation by nucleotide-sugar interconversion pathways. Eleven gene families have been identified that encode enzymes of the nucleotide-sugar interconversion pathways involved in the formation of the basic sugar building blocks of many plant cell walls (Penning et al. 2009; Yin et al. 2011). Although a number of different gene family members are regulated especially in response to invading *F. graminearum* hyphae and *T. dactyloides* pollen tubes (Figure 2E), there is almost no overlap in regulated genes, except for the *UAM5d* gene encoding a RGP (Reversibly Glycosylated Protein) which is downregulated both, in response to *F. graminearum* and *U. maydis*. Altogether, these first findings indicate already highly specific responses towards the various invaders.

### Incompatible pollination triggers among others downregulation of rehydration and cell wall-associated genes

To explore the molecular mechanisms underlying compatible and incompatible pollen-silk interactions, respectively, we compared the transcriptional responses of maize silks after 8 hours of pollination with B73 pollen (compatible) and *T. dactyloides* pollen (incompatible). To capture only maize silks’ responses, we used transcriptomic samples of maize pollen tubes to subtract the genes expressed in the male component. Notably, incompatible pollination resulted in a significant larger specific transcriptomic response in maize silks (32 versus 2,232 genes; Figure 3A). Notably, the expression pattern of the 113 common differently expressed genes (Figure 3A) after compatible and incompatible pollination was exactly the same (Figure 3B, Supplemental Data set 2). Out of the 113 genes, only one gene (*LHY1* - *late hypocotyl elongation protein ortholog1*) was downregulated in response to both conditions. The other 112 genes were upregulated (Figure 3B). These results suggest that the common genes are important for recognizing the presence of pollen grains but are not necessarily involved in the differentiation between compatible and incompatible pollen.

**Figure 3.**
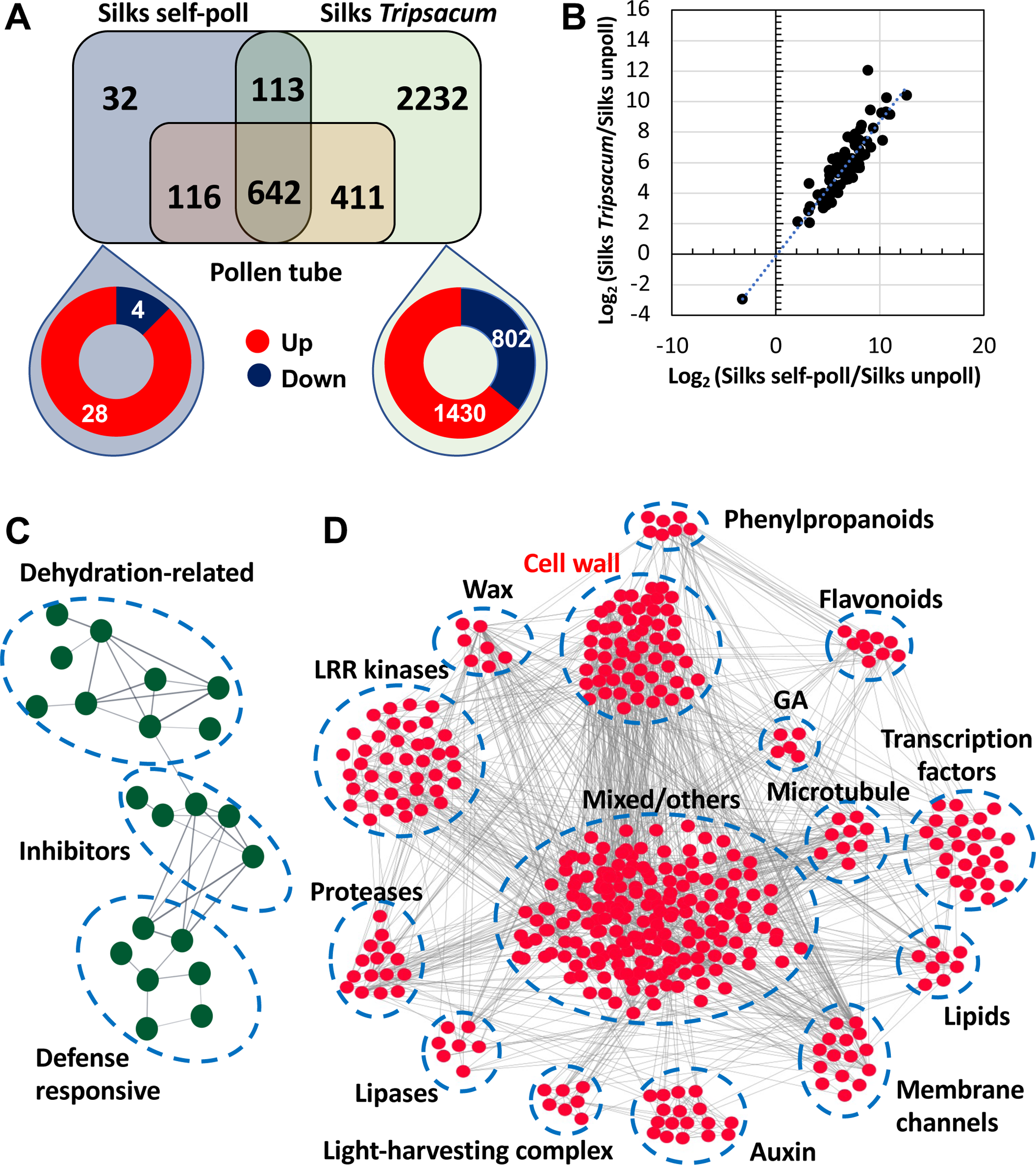
Comparison of transcriptional response in maize silks after compatible and incompatible pollination. **(A)** Comparison of differentially expressed genes (DEGs) between self-pollinated (compatible) and *Tripsacum dactyloides* pollinated (incompatible) silks. The pollen tube transcriptome was included as a control to identify silk-responsive genes to pollination. **(B)** Scatter plot representing log_2_ (fold change expression) for common DEGs in self-pollinated maize (x-axis) *versus* maize pollinated with *T. dactyloides* (y-axis). Network analysis of down-regulated DEGs after **(C)** compatible and **(D)** incompatible pollination of maize silks, respectively.

The strong transcriptional response in maize silks elicited by *T. dactyloides* pollen tubes (2,232 regulated genes) showed that about 36% genes were downregulated, which indicates a strategy developed by maize silks to shut down key pathways for pollen tube growth support ultimately leading to arrest of alien pollen tubes (Supplemental Data set 3). We then asked, which biological processes were downregulated by *T. dactyloides* pollination and which processes are important for successful compatible pollen tube growth. We conducted a cluster correlation analysis using the “cor” function implemented in the R package WGCNA to identify groups of genes with shared expression profiles. Since the maize genome is not well annotated and functions of many genes are categorized as unknown, it was likely that our gene network correlation analysis would contain a large portion of uncharacterized genes. Therefore, we combined our gene cluster correlation analysis with the web-tool STRING (Szklarczyk et al. 2015), which provides protein interaction data to be able to extract pathways that are affected by incompatible pollination. For compatible pollination we identified genes associated with dehydration-related, inhibitors and defense responsive as biological processes associated to compatible pollination (Figure 3C).

For incompatible pollination, our differential expression analysis yielded 803 downregulated genes. After removal of unrooted genes (384), the remaining downregulated differentially expressed genes were queried in STRING, which identified 419 interacting genes associated with incompatible pollination. Using a high edge confidence of at least 0.7, we constructed a gene network analysis that functionally grouped down-regulated genes into fifteen main hubs (Figure 3D, Supplemental Data Set 4). The largest group is formed by genes with a variety of functions, but the number of genes were not enough to form a clear cluster. The remaining fourteen clusters belong to genes involved in phenylpropanoid, flavonoids, cell wall and wax biosynthesis, but encode also microtubule components, transcription factors, lipids, membrane channels, light-harvesting complex components, lipases and lipid biosynthesis enzymes, proteases, and kinases including leucine-rich repeat (LLR) kinases. Gibberellin and auxin biosynthesis genes were also identified. Plant hormones appear to play roles during pollen-pistil interactions in maize, but their detailed functions are unclear. For instance, ethylene and auxin have been shown to rapidly increase in maize silks after compatible pollination (Mól et al. 2004). Increased levels in auxin followed by a spike of gibberellic acid (GA) biosynthesis were also reported after fertilization (Dorcey et al. 2009). In our study, incompatible pollination with *T. dactyloides* induced a strong downregulation of auxin biosynthesis and response genes (Figure 3D), but not of ethylene genes. Decrease in transcriptional expression of theses major growth hormones points towards a switch from development and growth to defense responses.

Water transport from stigmatic pistil cells to desiccated pollen grains is essential for pollen germination (Firon et al., 2012). Noteworthy, four clusters involved in pollen rehydration and germination were downregulated by *T. dactyloides* pollination indicating recognition of alien pollen and activation of mechanisms to prevent invasion. These genes that encode, for example, enzymes for lipid and wax biosynthesis, specialized water channels or aquaporins, and leucine-rich repeat (LLR) kinases have been shown previously to play a positive role in regulating pollen rehydration and initial stages of pollen tube growth in the female reproductive tract (Lee and Goring 2021; Windari et al. 2021; Abhinandan et al. 2022). We found genes of several members of the aquaporin gene family that are downregulated after *T. dactyloides* pollination. In Arabidopsis, it has been shown that *SIP1;1* and *PIP1;2*, orthologous of the downregulated maize genes found in this study, are key regulators of water transport from papilla cells to pollen and thereby regulating pollen hydration (Windari et al. 2021). Similarly, in solanaceous and crucifer, but also lily plant species, pistil lipids and lipid transfer-like protein are required for pollen hydration and further penetration (Park et al. 2000; Sanchez et al. 2004; Thompson and Raizada 2018; Zheng et al. 2018). Furthermore, loss-of-function mutations in *RKF1*, an *LLR* kinases gene, reduced pollen hydration of wildtype pollen on mutant stigma (Lee and Goring 2021) indicating the importance of pollen recognition processes. In conclusion, although downregulation of LLR kinase, lipid, wax, and aquaporin genes after incompatible pollination are not sufficient to prevent initial germination and penetration of fast-growing *T. dactyloides* pollen tubes, they appear to be sufficient with other processes (see below) to prevent most incompatible pollen tubes to reach the ovaries.

The largest cluster of regulated genes is associated with the cell wall (Figure 3D, Supplemental Data Set 4). There were further sets of downregulated and related gene clusters formed by phenylpropanoid, flavonoids, and microtubule genes (Figure 3D). Phenylpropanoids are a structurally diverse group of phenylalanine-derived metabolites involved in the synthesis of flavonoids and lignins used by plants to build cell walls. Flavonoids have been well studied in pollen grains as they are plant-specific compounds required for pollen germination and tube growth, and are constituents of the pollen cell wall and work as positive effectors for pollen tube elongation (Muhlemann et al. 2018; Dong et al. 2021; Wu et al. 2023; Xue et al. 2023). Thus, downregulation of phenylpropanoids and consequently flavonoids likely arrests pollen tube growth as observed in incompatible pollinations of maize (Lausser et al. 2010). Similarly, cell wall and microtubule genes have been largely known for their importance during pollen tube/pistil interaction and pollen tube guidance (Dresselhaus and Franklin-Tong 2013; Riglet et al. 2020). During compatible pollination, these dynamic interactions facilitate pollen tube elongation. In plants of the Liliaceae family as well as in Arabidopsis, such interactions are mediated, for example, by microtubules, pectins, lipids and a stylar cysteine-rich adhesin protein secreted in the secreted extracellular matrix of the pistils (Park et al. 2000; Onelli et al. 2015; Riglet et al. 2020). In summary, our data shows downregulation of rehydration genes, microtubule and cell wall-related genes explaining pollen tube growth arrest in incompatible pollinations.

### Compatible pollination regulates especially expression of MYB transcription factor and senescence genes

Notably, as reported above, transcriptional responses in maize silks to compatible and incompatible pollination show only small overlap (Figure 3A). Intriguingly, compatible pollination resulted only in 32 differentially expressed unique genes (Supplemental Table S2). We found that 10 of these genes encode transcription factors (Supplemental Table S2). Genes involved in transcriptional regulation are likely to be important components of pollen-pistil interaction, since they can precisely regulate large numbers of downstream genes. Especially MYB transcription factor genes (15% of the total differentially expressed unique genes) were regulated during compatible pollination. In Arabidopsis, MYB transcription factors have been reported as key regulators of pollen tube growth and sperm release (Leydon et al. 2013), pollen tube-synergid interaction (Liang et al. 2013) and senescence (Cao et al. 2023). In maize, MYB transcription factor genes were reported as targets of high temperature leading to pollen sterility (Begcy et al. 2019). A whole genome analysis of gene expression profiles during pollen-pistil interactions in Arabidopsis showed that 16% of the differentially expressed transcription factor genes during the programic phase in pistils belongs to the *MYB* gene family. *AtMYB70* and *AtMYB123* were identified as major regulators of two of the differentially expressed clusters (Boavida et al. 2011). Functional studies are now necessary to evaluate the MYB transcription factor identified in this study and to elucidate their contribution to compatible pollen-pistil interaction(s) in maize.

Previous studies have reported that maize silks undergo an accelerated process of senescence in response to compatible pollination (Bassetti and Westgate 1993; Sella Kapu and Cosgrove 2010; Šimášková, et al. 2022). Thus, we searched for senescence-associated genes within the differentially expressed genes in maize silks as a result of compatible pollination. We found that 34% of these genes are indeed linked to senescence previously reported in maize and lettuce leaves (He et al. 2020; Chase et al. 2024). Thus, our data indicate that silk and leaf senescence could be controlled by similar mechanisms. Senescence is a complex biological process controlled by multiple genetic and environmental variables. Many associated senescence genes have been identified and transcription factors have been recognized as hub genes in senescence pathways in many plant species (Bengoa Luoni et al. 2019; Cao et al. 2023). We found four transcription factor genes in the list of DEGs in maize silks after self-pollination (Supplemental Table S2) previously associated with senescence (Figure 4B). The homolog of *ZmMYBR79*, also known as *CIRCADIAN CLOCK-ASSOCIATED 1* (*CCA1*) was previously shown to inhibit leaf senescence. *mybr/cca* mutants showed accelerated age-dependent senescence (Song et al. 2018). Notably, *ZmMYBR79/CCA1* was downregulated 3.85-folds after compatible pollination. Since *ZmMYBR79/CCA1* forms a core loop with *LATE ELONGATED HYPOCOTYL1* (*LHY1*) by repressing one another’s expression (Song et al. 2018), we also studied LHY transcriptional expression after compatible pollination and found that *ZmLHY* was also downregulated (Figure 4B). Similarly, *ZmMYBR48*, another gene that has been associated with the circadian clock (Huang et al. 2017) and which is likely also contributes to the *ZmMYBR79/CCA1-ZmLHY1* senescence regulatory network, was found downregulated after compatible pollination (Figure 4B). Our data suggests that silk senescence in maize is likely accelerated due to downregulation of a *ZmMYBR79/CCA1-ZmLHY1-ZmMYBR48* regulatory network. Mutant studies are now required to support this assumption.

**Figure 4.**
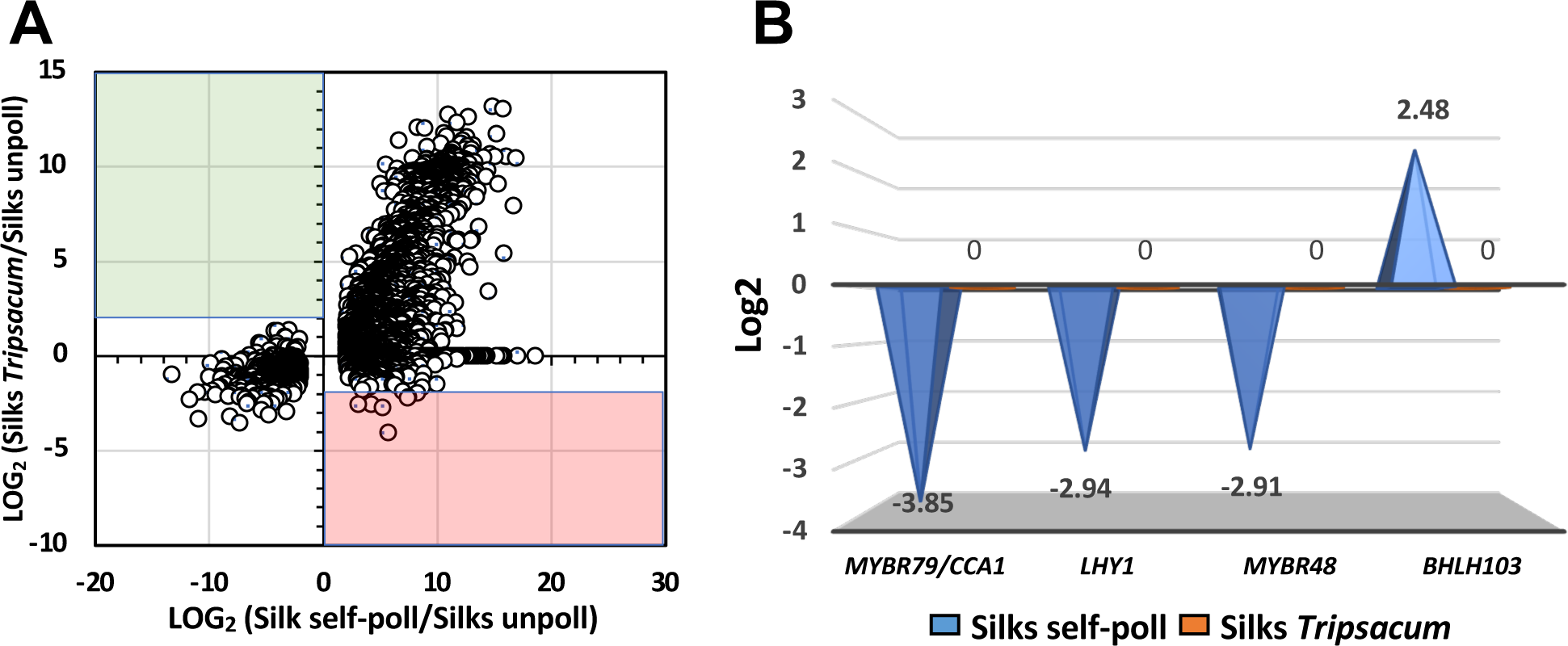
Self-pollination alters expression of senescence-associated transcription factor genes in maize silks. **(A)** Self and incompatible pollination induce large transcriptional expression as showed in the scatter plot representing the log_2_ (fold change expression) for all DEGs in self-pollinated maize (x-axis) *versus* maize silks pollinated with *T. dactyloides* (y-axis). **(B)** Fold change expression changes of senescence associated transcription factor genes.

Another transcription factor differentially expressed after compatible pollination is *bHLH103* (FC=2.48, Figure 4B). Basic helix-loop-helix (bHLH) transcription factors play important roles in regulating multiple biological processes in plants. In petunia, for example, a bHLH transcription factor (FBH4) was shown to regulate flower senescence. While overexpression of *FBH4* resulted in accelerated flower senescence, RNAi plants showed retarded senescence (Yin et al. 2015). Therefore, the coordinated downregulation of MYBs as senescence repressors and upregulation of bHLHs as likely senescence promoters indicates that precocious silk senescence is initiated after compatible pollination likely to prevent growth of other biotic invaders.

### Downregulation of cellulose biosynthesis genes occurs during incompatible pollination and fungal invasion

Rigidification of the silks’ cell wall has been reported as one of the consequences in response to pollination in maize (Sella Kapu and Cosgrove 2010). Similarly, host cell wall damage has been observed during pathogen infection (Lorrai and Ferrari 2021). Therefore, we next investigated cell wall-related responses to incompatible pollination and fungal invasion (Figure 5). We focused our strategy on gene families whose products function in six stages of cell wall biogenesis: substrate generation, polysaccharide synthesis, membrane trafficking, assembling and turnover, secondary wall formation, and signaling (Penning et al. 2009). Cell wall biosynthesis begins with the conversion of sucrose into glucose and fructose molecules for conversion into storage carbohydrates by sucrose synthases genes (Stein and Granot 2019). Thus, we also checked the expression of these genes. Cellulose is a major constituent of plant cell walls. It is synthesized at the plasma membrane by the cellulose synthase complex (CSC) and deposited as cellulose polymers into the developing cell wall (Watanabe et al. 2018). We first explored the transcriptional expression of cellulose synthase (*Ces*) and cellulose synthase-like (*Csl*) genes (Figure 5A, B). In maize, the *Ces* family consists of 20 members. Incompatible pollination reduced the expression of cellulose synthase (CesA) genes, particularly that of *CesA1*, *CesA2*, *CesA4* and *Ces7b* (Figure 5A). Even though *F. graminearum* and *U. maydis* invasion downregulated members of the cellulose synthase gene family, these group of genes were different than the ones downregulated by *T. dactyloides*. Both fungal pathogens reduced the transcriptional expression of *CesA11a*, *CesA11b* and *CesAL4*, respectively. Notably, an EMS-induced mutant, *brittle stalk-5* (*BK-5*), which encodes a cellulose synthase (*CesA11b*) involved in cellulose biosynthesis was characterized using genetic analysis and map-based cloning (Li et al. 2022). Infection by both fungal pathogens repressed *CesA11b* by 9,2-fold and 4.4-fold, respectively. No changes in expression were found during incompatible pollination (Figure 5A). Since *CesA11b* is critical for cell stiffness, infection by F. *graminearum* and *U. maydis* might be associated with silk brittleness as a strategy to prevent fungal progression.

**Figure 5.**
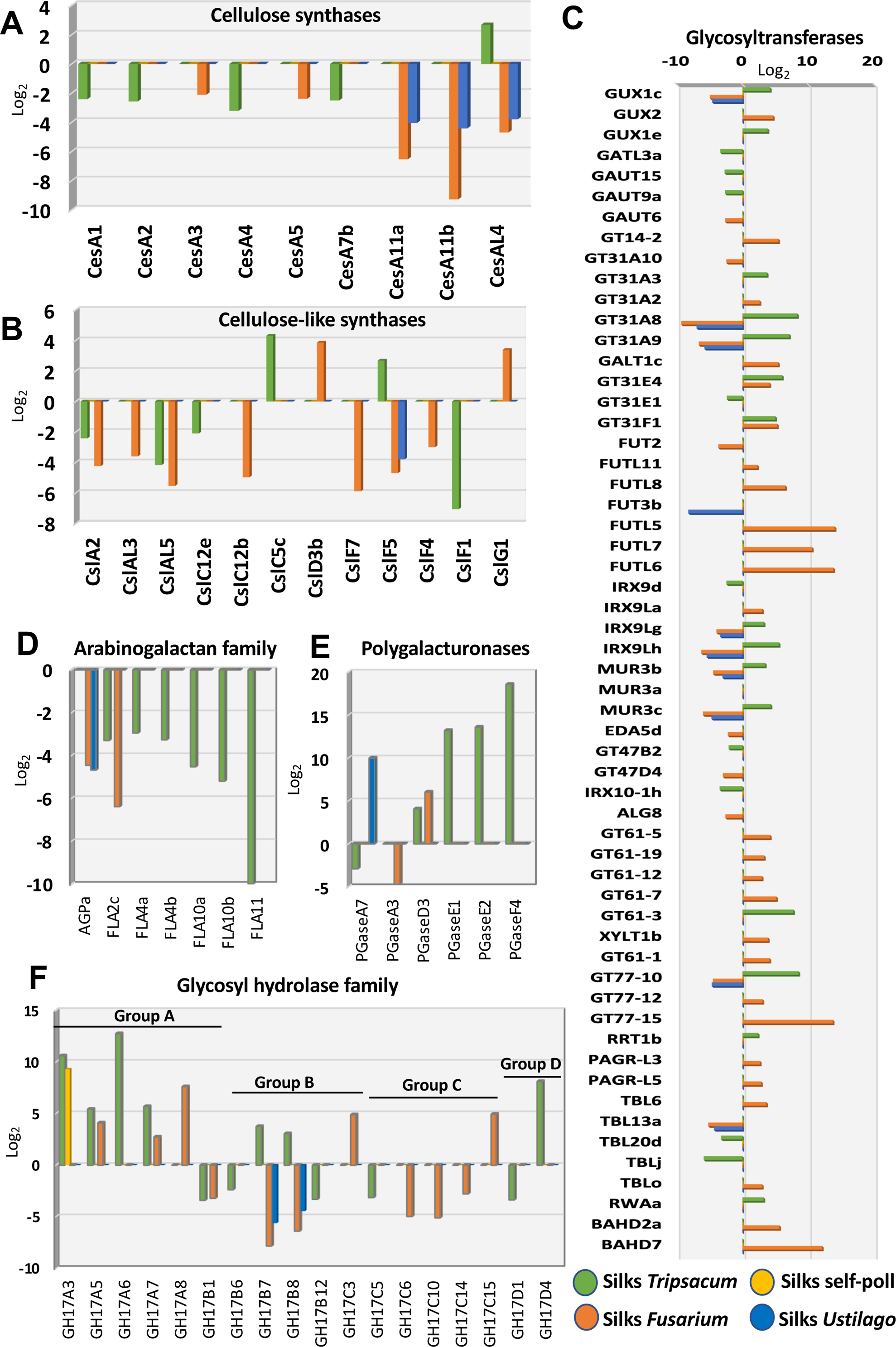
In contrast to self-pollination, incompatible pollination and fungal invasion alters cell wall-related genes families in maize silks. Fold change expression analysis of gene families related to cell wall formation are shown. Genes encoding cellulose synthases **(A)** and most genes for cellulose-like synthases **(B)** are down-regulated. Genes encoding glycosyltransferases **(C)** are differentially expressed, while genes involved in arabinogalactans **(D)** are down-regulated and polygalacturonanases **(E)** are mostly up-regulated. Genes for glycosyl hydrolases **(F)** show a differential expression pattern.

In addition to *CesA* genes, maize also has cellulose synthase-like (*Csl*) genes that synthesize other essential non-cellulosic polysaccharides that are constituents of the cell wall (Li et al. 2019). Genes belonging to the *Csl* gene family were also downregulated after invasion of alien pollen tubes and fungal hyphae (Figure 5B). While pollination by *T. dactyloides* and pathogen infection by *F. graminearum* led to a larger number of downregulated *Csl* genes, *CslF5* was the only gene with reduced expression after *U. maydis* infection in maize silks. It is noteworthy that pollination by compatible maize pollen (self-pollination) did not trigger any transcriptional expression on *CesA* and *Csl* genes, respectively. In summary, these results indicate that cellulose biosynthesis is downregulated when non-compatible pollen or fungal pathogen interact and grow within maize silks likely contributing to fast growth inhibition.

### Glycosyltransferase and -hydrolase genes are strongly regulated in response to alien invaders

Another important super gene family primarily involved in cell wall formation encodes glycosyltransferases. These are transmembrane proteins localized in the endoplasmic reticulum and Golgi apparatus of the plant secretory system (Amos and Mohnen 2019). Cell wall glycosyltransferases are involved in several processes including synthesis of the polymer backbones, elongation of side branches with extended glycosyl chains, and addition of single monosaccharide linkages onto polysaccharide backbones and/or side branches (Amos and Mohnen 2019). The glycosyltransferases are divided into processive and non-processive enzymes. Within the non-processive glycosyltransferase gene families, twelve families showed differential responses to incompatible pollination and fungal invasion but remained unaltered after self-pollination (Figure 5C). Glycosyltransferases are involved in pectin synthesis (Qin et al. 2013; Zabotina et al. 2021). Glucuronic acid substitution of xylan (*GUX1c*) was the only gene differentially expressed by incompatible pollination and fungal invasion (Figure 6C). Notably, *GUX1c* was downregulated after infection by both F. *graminearum* and *U. maydis* and upregulated by *T. dactyloides*. *GUX1c* did not change in response to compatible pollination. In Arabidopsis, *gux1/2/3* triple mutants cause a reduction in plant growth and in the mechanical strength of the stem (Lee et al. 2012).

**Figure 6.**
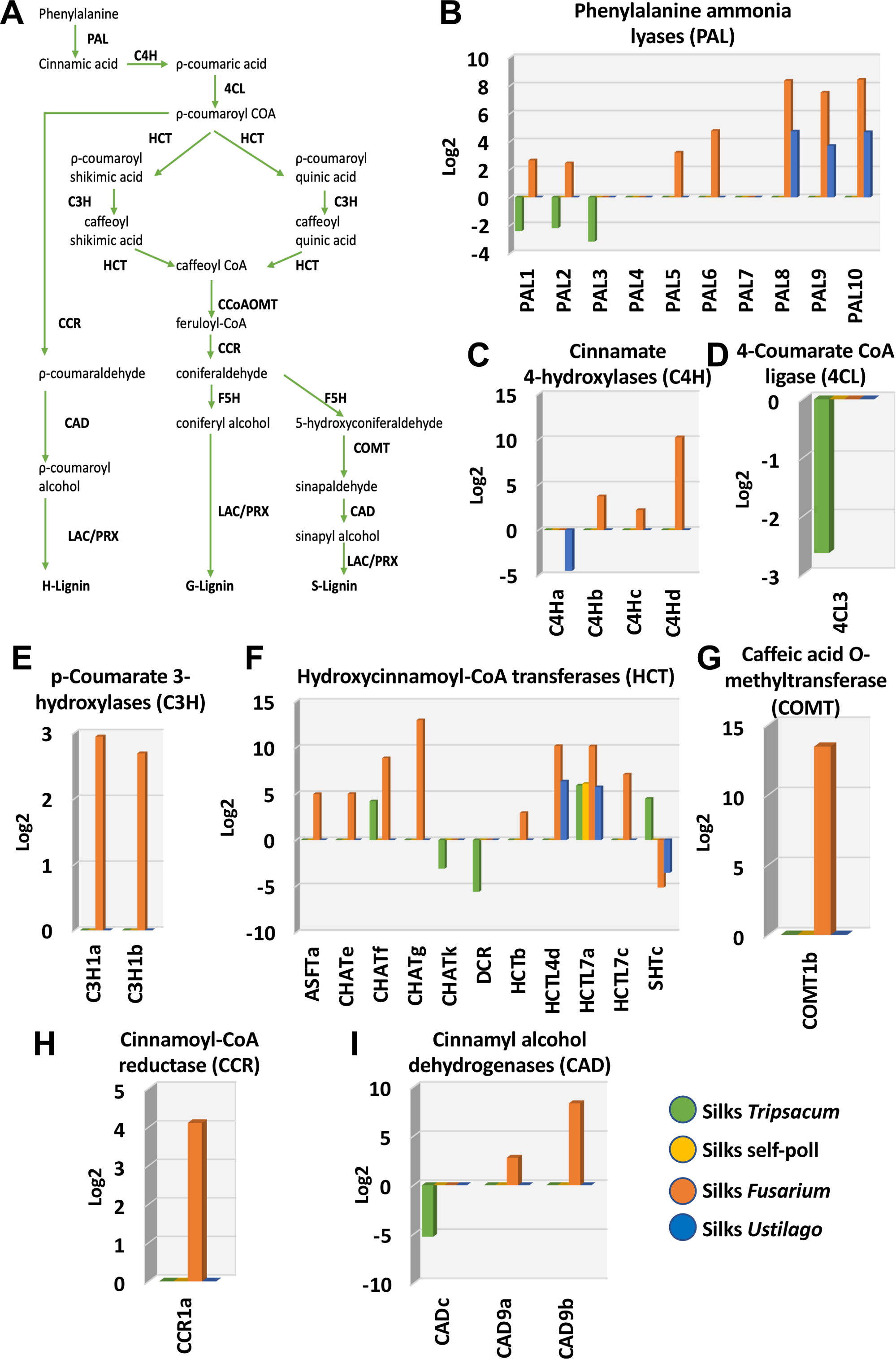
Genes involved in monolignol biosynthesis are highly upregulated after *F. graminearum* infection in maize silks. **(A)** Overview of the H-/G-/S-monolignol biosynthesis pathway including products, educts, and respective enzymes. **(B-I)** Expression pattern of monolignol biosynthesis genes in maize silks including genes encoding (B) phenylalanine ammonia lyases (PAL), (C) cinnamate 4-hydroxylases (C4H), (D) 4-coumarate CoA ligases (4CL), (E) p-coumarate 3-hydroxylases (C3H), (F) hydroxycinnamoyl-CoA transferases (HCT), (G) caffeic acid O-methyltransferases (COMT), (H) cinnamoyl-CoA reductases (CCR) and for (I) cinnamoyl alcohol dehydrogenases (CAD).

Functions of the *GT31* family is associated with N-glycan formation, however, it also works in galactan backbone formation of arabinogalactan-proteins (Showalter and Basu 2016; Nibbering et al. 2022). We found strong upregulation of *GT31A8* and *GT31A9* in response to incompatible pollination and downregulation in response to F. *graminearum* and *U. maydis*. *GT31A10* was downregulated, but only in response to F. *graminearum*. All remaining members differentially expressed within the GT31 family were upregulated in response to *T. dactyloides* and *F. graminearum* (Figure 5C). *GT43* and *GT47* gene families were downregulated in response to *F. graminearum* and *U. maydis* and upregulated in response to *T. dactyloides*. *GT43* is predicted to contain genes encoding the (1→4)-β-d-xylan synthase of glucuronoarabinoxylans (Mitchell et al. 2007) involved in the biosynthesis of xylose, pectin and xyloglucan (Wu et al., 2019). Similarly, the *GT47* gene family was upregulated in response to incompatible pollination and downregulated in response to fungal invasion (Figure 5C). *GT61*, *GT77* and *GT106* gene families were mostly upregulated in response to incompatible pollination and fungal invasion (Figure 5C). While *GT61* is involved in the synthesis of cell wall xylans (Cenci et al. 2018), *GT77* and *GT106* families function has been associated to pectin biosynthesis (Møller et al. 2017; Takenaka et al. 2018).

Arabinogalactan proteins constitute a large highly glycosylated family of hydroxyproline-rich proteins involved in plant reproduction. Notably, both incompatible pollination and fungal invasion downregulate arabinogalactan genes (Figure 5D). The strongest impact on this gene family was observed in response to *T. dactyloides* pollination. Six arabinogalactan protein members were downregulated. In tomato and Arabidopsis, expression of arabinogalactan genes has been observed particularly during the progamic phase in the pistil suggesting a potential function during pollen–pistil interaction (Pereira et al. 2014; Lara-Mondragón and MacAlister 2021). Since this gene family was predominately affected by incompatible pollination, they could be a good target for future incompatibility studies during pollination in maize.

Similarly, as observed with arabinogalactans, glycosyl hydrolases including polygalacturonases (pectinases) were differentially expressed in response to *T. dactyloides* (Figure 5E, F). Pectinases are enzymes that cleave pectin, a major cell wall component especially accumulated in the middle lamellae between plant cells and which have been shown to participate in cell wall elongation and pollen tube growth (Minic 2008; Babu and Bayer 2014; Lu et al. 2021). We found that incompatible pollination enhanced expression of polygalacturonase genes (Figure 5E). This suggests that *T. dactyloides* softens cell walls, opposite of what it is observed after compatible pollination, where cell wall undergoes structural modifications leading to its rigidification (Sella Kapu and Cosgrove 2010). Among other glycosyl hydrolases, *U. maydis* infection only downregulated *GH14B7* and *GH14B8* genes (Figure 5F). *F. graminearum* and *T. dactyloides* induced mixed responses. While glycosyl hydrolases from group A were upregulated, group C were downregulated. Groups B and D showed both pattern of expression (Figure 5F). In summary, many genes encoding glycosyltransferases and hydrolases are regulated in response to incompatible pollination and fungal invasion likely leading to cell wall modifications that suppress fast growth and cell elongation of alien invaders in silk tissue.

### *Fusarium graminearum* triggers the monolignol biosynthesis pathway in maize silks

Biosynthesis of lignin, one of the core components of cell walls that form a complex network of polymers of hydroxylated and methoxylated phenylpropane units, depends on the phenylpropanoid and monolignol pathways. Both pathways are interconnected since monolignol biosynthesis utilizes substrates generated by the phenylpropanoid pathway (Yadav et al. 2020). We found that *F. graminearum* induced high expression of key genes involved in the monolignol biosynthesis pathway (Figure 6A) in maize silks. The conversion activity of phenylalanine ammonia lyases (PALs) into cinnamic acid is the first step of the general phenylpropanoid pathway. Seven maize *PALs* showed significant upregulation compared with both mock and untreated samples, particularly *PAL8*, *PAL9* and *PAL10*, which showed eight-time fold increase (Figure 6B). Similarly, the three genes showed high expression in response to *U. maydis*. In contrast, *T. dactyloides* pollination decreased *PAL1*, *PAL2* and *PAL3* expression by more than 2-fold. *PAL* genes were described as mediators for cell wall’s first layer of immunity and involved in broad spectrum disease resistance (Ning and Wang 2018; Zhou et al. 2018). Thus, upregulation of *PAL* genes after fungal invasion is a mechanism of defense also used by maize silks to prevent the spread of pathogens. In contrast, incompatible pollination does not activate this immune response.

Cinnamate 4-hydroxylase (C4H), involved in the reduction of cinnamic acid into ρ-coumaric acid, showed also high upregulation in response to *F. graminearum* (Figure 6C). *C4Hb*, *C4Hc*, and *C4Hd* showed a 3.8-fold, 2.2-fold, and 10.4-fold increase, respectively. Compatible and incompatible pollination did not alter any of the *C4H* genes. *U. maydis* infection led to downregulation of *C4Ha* but did not alter the expression of any other member of this gene family (Figure 6C). Notably, neither pathogen infection nor compatible pollination changed 4-coumarate CoA ligase (*4CL*) expression. Only *T. dactyloides* pollination triggered decreased *4CL* expression (Figure 6D).

An important step in the general phenylpropanoid pathway involves the conversion of ρ-coumaroyl-CoA to caffeoyl-CoA by the p-coumarate3-hydroxylase (*C3H*) and hydroxycinnamoyl-CoA:shikimate/quinate hydroxycinnamoyltransferase (*HCT*). Gene expression changes of *C3H* by compatible and incompatible pollination as well as *U. maydis* infection were not observed. Only *F. graminearum* infection induced the expression of both sets of genes in maize silks. While *C3H1a* and *C3H1b* were induced 2.95-fold and 2.68-fold compared to the mock treatment (Figure 6E), respectively, nine *HCT*s were differentially expressed (Figure 6F). Eight of them were upregulated and one downregulated (*sHTc*), respectively. Similarly, *U. maydis* invasion upregulated *HCTL4d* and *HCTL7a* in maize silks and downregulated *SHTc*. Incompatible pollination by *T. dactyloides* downregulated *CHATk* and *DRC*, and upregulated *CHATf*, *HCTL7a*, and *SHTc*. Compatible pollination upregulated *HCTL7a* similarly as incompatible and fungal invasion (Figure 6F).

Notably, Arabidopsis leaves challenged with several pathogens (Zhang et al. 2018; Vogel et al. 2021; Zhu et al. 2023; Entila et al. 2024) induced the expression of the caffeic acid (5-hydroxyconiferaldehyde) O-methyltransferase gene (*COMT*1), similarly as we observed in maize silks infected with *F. graminearum* (Figure 6G). Regulation of *COMT* was suggested to be one of the first mechanisms of plant’s pathogen defense and obviously occurs also in maize silks.

The general phenylpropanoid and monolignol pathways share genes involved in the initial biosynthesis steps (Figure 6A). *F. graminearum* not only induced genes involved in the general phenylpropanoid pathway but also in the monolignol pathway. Sinapyl alcohol and coniferyl monolignol are the major building blocks of lignin and they generate syringyl (S) and guaiacyl (G) units of lignin polymers, respectively. The minor monolignol unit, which is deposited more in monocots, is the p-hydroxyphenyl (H) unit derived from p-hydroxyphenyl (Yadav et al. 2020). Cinnamoyl-CoA reductase (*CCR*) converts five types of hydroxycinnamoyl-CoA into cinnamaldehyde and their overexpression was shown to play roles in pathogen defense responses of plants (Lauvergeat et al. 2001; Cheng et al. 2017). Only *F. graminearum* infection induced *CCR1a* in maize silks (Figure 6H). Additionally, *F. graminearum* infection upregulated *CAD9a* and *CAD9b* in maize silks (Figure 6I). Cinnamyl alcohol dehydrogenases (*CADs*), which deoxidizes five types of cinnamaldehyde to the corresponding monolignols (Figure 6A) were also associated with lignin synthesis and stress resistance (Guo et al. 2010).

Peroxidases (PRX) and laccases (LAC) are further classes of enzymes present in the apoplast of plant cells. Among their diverse functions, they are also important for lignin biosynthesis (Laitinen et al. 2017). In our study, forty genes encoding PRX enzymes were regulated in all conditions: 18 after *T. dactyloides* pollination, 27 after *F. graminearum* and only three after U. *maydis* infection (Figure 7A). Only one gene, *PRDG7* was upregulated after compatible pollination. Noteworthy, the same gene was upregulated after *T. dactyloides* pollination, which suggests that it responds to pollination in general. Remarkably, the pattern of expression of *PRX*s is opposite when incompatible pollination and fungal invasion by *F. graminearum* is compared. While *T. dactyloides* pollination represses *PRX*s, *F. graminearum* infection enhances *PRX*s expression. Another gene family involved in the production of lignin is *LAC*. After *F. graminearum* infection, *Lac5b, Lac5c, Lac7a, Lac7c, Lac7k, Lac15a*, and *Lac15b* showed increased expression, while *Lac7h* was downregulated (Figure 7B). Only *Lac7a* increased in response to U. *maydis* and *Lac15a* decreased after *T. dactyloides* pollination. None of the laccases responded after compatible pollination. These results show that incompatible pollination negatively impacts the entire phenylpropanoid and monolignol pathways, while fungal infection promotes the expression of genes involved in the lignin biosynthesis pathway.

**Figure 7.**
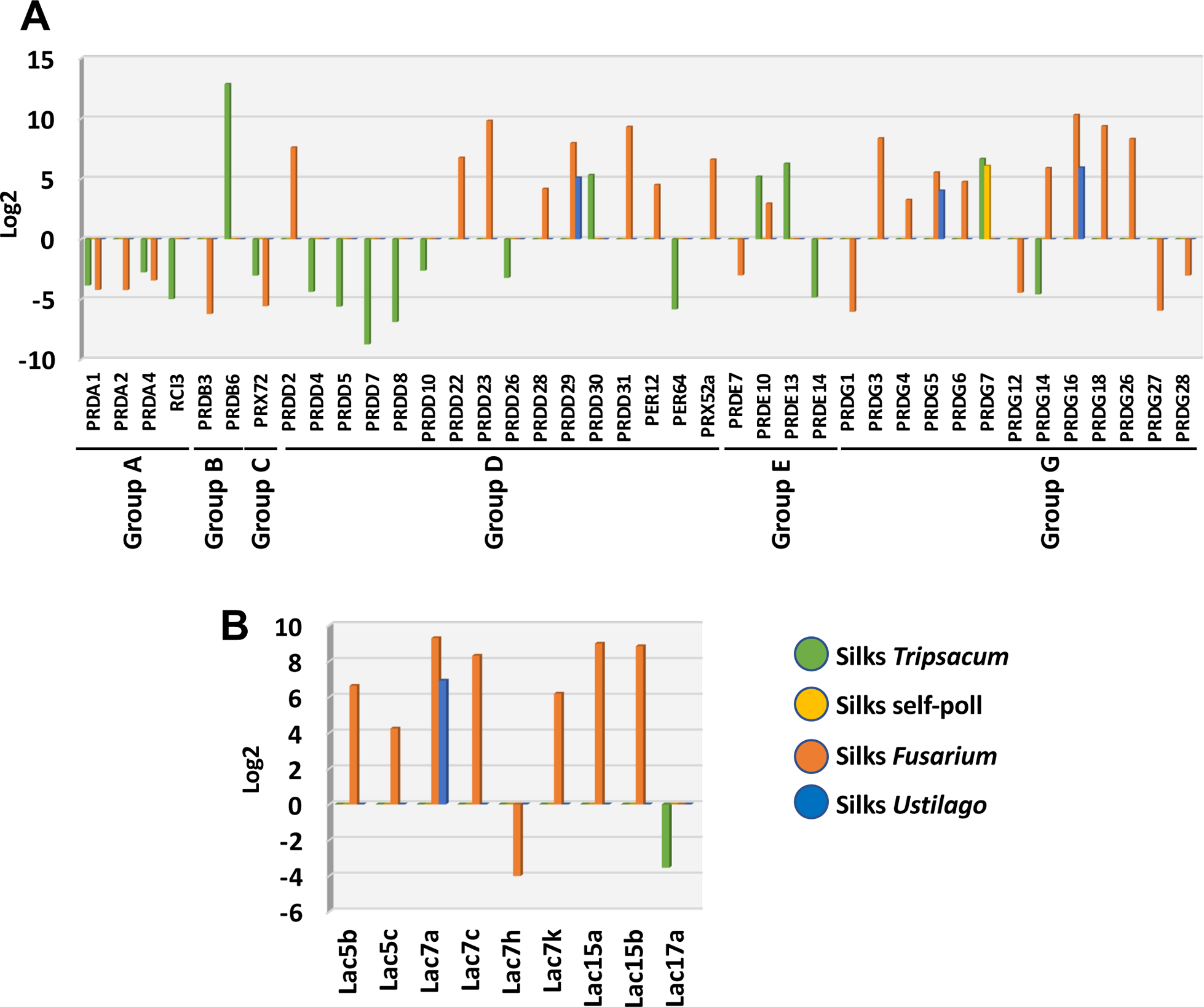
Genes encoding enzymes that facilitate ROS reduction and lignin polymerization are only regulated in maize silks by foreign invaders. **(A)** Peroxidase (PRX) and **(B)** laccase (LAC) genes are largely up-regulated after *F. graminearum* infection, partly up-regulated after *U. maydis* infection and tendentially down-regulated by pollen tubes of *T. dactyloides*.

### NAC and WRKY transcriptional regulators are highly upregulated in response to fungal infection, but not to pollination

Activation of monolignol biosynthesis pathways and polymerization of the monolignol units is genetically controlled by an evolutionary conserved regulatory network of NAC and MYB transcription factors (Ohtani and Demura 2019). To elucidate whether the high expression of the monolignol pathway after *F. graminearum* infection correlates with a NAC-MYB gene regulatory network in maize silks, we next studied the expression of the two gene families. Four secondary wall-associated NACs (*ZmSWN1, ZmSWN3, ZmSWN6* and *ZmSWN7*) have been described as members of the NAC-MYB gene regulatory network (Zhong et al. 2011). None of these genes is expressed in maize silks neither in treated nor in control samples. However, they are expressed in root tissues and *ZmSWN7* is also expressed in leaves (Supplemental Figure S3). Since the characterized secondary wall-associated *NAC* genes were not expressed in maize silks, we examined the entire NAC transcription gene family to identify putative NAC regulators in maize silks. In comparison to vegetative tissues like roots and leaves, expression of *NAC* genes in unpollinated mature silks was low (Figure 8A). Expression of *NAC* genes increased during silk aging but was especially high after fungal infection (Figure 8A, B). For instance, while *ZmNAC109*, *ZmNAC75*, and *ZmNAC5* were particularly induced by *F. graminearum*, *ZmNAC23*, *ZmNAC30* and *ZmNAC4* were upregulated by both, *F. graminearum* and *U. maydis* infection (Figure 8B). These differences could explain why many more genes are regulated in maize silks after *F. graminearum* compared with *U. maydis* infection.

**Figure 8.**
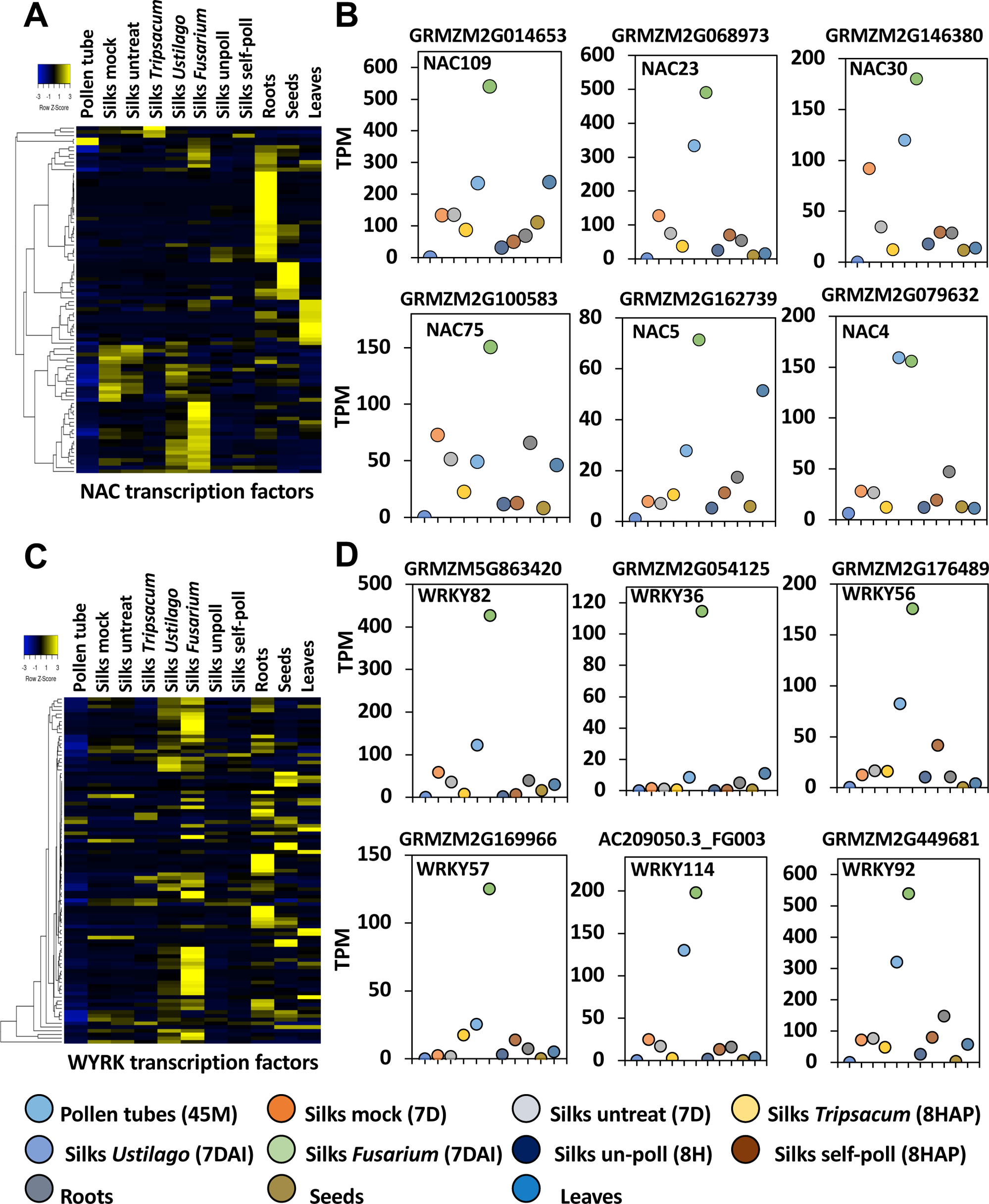
NAC and especially WRKY transcription factor genes are induced after fungal infection in maize silks. **(A)** Heat map displaying induction of NAC transcription factor genes in maize silks at indicated conditions. **(B)** Transcript per million reads (TPM) of selected NAC transcription factor genes. **(C)** Heat map displaying induction of WYRK transcription factor genes in maize silks at indicated conditions. **(D)** TPMs of selected WYRK transcription factor genes indicate high induction especially after *F. graminearum* infection.

Some MYB transcription factors function downstream of NACs as a second layer of transcriptional regulation (Ohtani and Demura 2019). Targets of these MYB proteins are among others enzyme-encoding genes involved in xylan and lignin biosynthesis. Thus, we next investigated expression of the entire maize MYB transcription factor family and their responses to pollination and fungal invasion (Supplemental Figure S4A). We found that, for example, *ZmMYB20* and *ZmMYB100* were highly upregulated after *T. dactyloides* pollination (Supplemental Figure S4B-C), but a strong correlation to specific invaders like above-described NAC transcription factor genes was not observed.

Another class of important gene regulators acting in plant defense mechanisms are WRKY transcription factors. So far, WRKYs have been comprehensively studied using mainly Arabidopsis as plant model organism (Seo et al. 2015). In maize silks, members of the *WRKY* gene family were highly induced in response to pathogen infection, particularly by *F. graminearum* (Figure 8C). Three genes, *ZmWRKY36*, *ZmWRKY57* and *ZmWRKY82* are almost specifically upregulated after *F. graminearum* infection, while *ZmWRKY56*, *ZmWRKY92* and *ZmWRKY114* are upregulated by both pathogens, but significantly higher by *F. graminearum* (Figure 8D). WRKY transcription factors have been shown to positively regulate jasmonate (JA)-mediated plant defense to necrotrophic fungal pathogens (Chen et al. 2021). Two JA-associated genes, octadecanoid-responsive Arabidopsis (*ORA59*) and plant defensin 1.2a (*PDF1.2*) were upregulated in *AtWRKY75* (a homolog of *ZmMWKY36*) overexpressing Arabidopsis transgenic plants (Chen et al. 2021). The maize ortholog of Arabidopsis *ORA59* has been annotated as *Ereb58* (AP2-EREBP-transcription factor 58). *F. graminearum* induced its expression 6.5-fold in maize silks. Expression of the maize ortholog of *PDF1.2* (defensin-like protein2, *Def2*), however, neither changed in response to pathogens nor to pollination (Supplemental Data Set 1). Further investigations are now necessary to identify targets of above-described NAC and WRKY transcription factors to better understand differences between the gene regulatory networks activated after *F. graminearum* and *U. maydis* infection and to develop tools to prevent growth and spread of both pathogens.

### *F. graminearum* and *U. maydis* provoke a diterpenoid response in maize silks

To further investigate the differential response of maize silks to *F. graminearum* and *U. maydis* infection, we next compared the global transcriptional response. Even though a large overlap of DEGs was detected (Figure 9A), *F. graminearum* induced a significantly larger differentially expressed response compared to *U. maydis* (Figure 9B). Gene ontology analysis showed enrichment of responses to stimuli, macromolecule and protein modification as main functional processes affected by *F. graminearum.* Divergently, *U. maydis* mostly impacted metabolic processes associated with photosynthesis (Supplemental Figure S5). Considering that these findings are very general, we searched for pathways that are associated with the processes and selected the terpenoid pathway as an example as it is involved as a major component of maize defenses to herbivores, pathogens, and other environmental challenges (Mafu et al. 2018). In maize, the antifungal phytoalexins kauralexins and dolabralexins, but also the hormone gibberellic acid (GA) derive from a common precursor, ent-CPP. Geranyl geranyl diphosphate (GGPP) is the precursor of ent-CPP (Figure 9C). Two catalytically redundant diterpene synthase (diTPS) enzymes, ANTHER EAR 1 (ZmAn1) and ANTHER EAR 2 (ZmAn2) control the production of ent-CPP (Schmelz et al. 2011; Mafu et al. 2018; Murphy et al. 2021). Infection by *F. graminearum* resulted in a more than 14-fold increase in *ZmAN2* expression (Figure 9D). *ZmAN1* and *ZmAN2* did not change in response to *U. maydis*. It has been shown that *ZmAN2* is critical for kauralexin and dolabralexin biosynthesis, while Z*mAN1* controls GA biosynthesis. This allows a pathway partition by separating precursor flux toward primary and secondary diterpenoid pathways, respectively (Mafu et al. 2018; Ding et al. 2019; Murphy et al. 2021). *ZmKSL2* (kaurene synthase-like2) and *ZmKSL4* drive the production of kauralexin and dolabralexin biosynthesis, respectively. Genes for both enzymes are upregulated in response to *F. graminearum* infection (Figure 9E, F), while only *ZmKSL2* is induced after *U. maydis* infection (Figure 9E). Cytochrome P450 monooxygenase 16/18 (ZmCYP71Z16/18), another enzyme required to generate kauralexin and dolabralexin, was induced 12-fold and 8-fold in response to *F. graminearum* and *U. maydis*, respectively (Figure 9G). Notably, the maize antifungal toxin zealexin is produced in an initially independent pathway using ZmdiTPS6/11 (diterpene synthases). *ZmdiTPS6/11* was only upregulated in response to *F. graminearum* (Figure 9H). In contrast to genes required for phytoalexin biosynthesis, *ZmKSL5* and *KO1* representing key genes for GA biosynthesis were downregulated after *F. graminearum* infection (Figure 9I, J). In summary, these results show that *F. graminearum* infection triggered the expression of phytoalexin biosynthesis genes, while genes involved in GA biosynthesis are downregulated. In contrast, only minor or no changes in terpenoid pathway genes was triggered after *U. maydis* infection. These findings are in agreement with a previous study showing that *F. graminearum* infection activates the generation of phytoalexins in root tissues of maize (Mafu et al. 2018).

**Figure 9.**
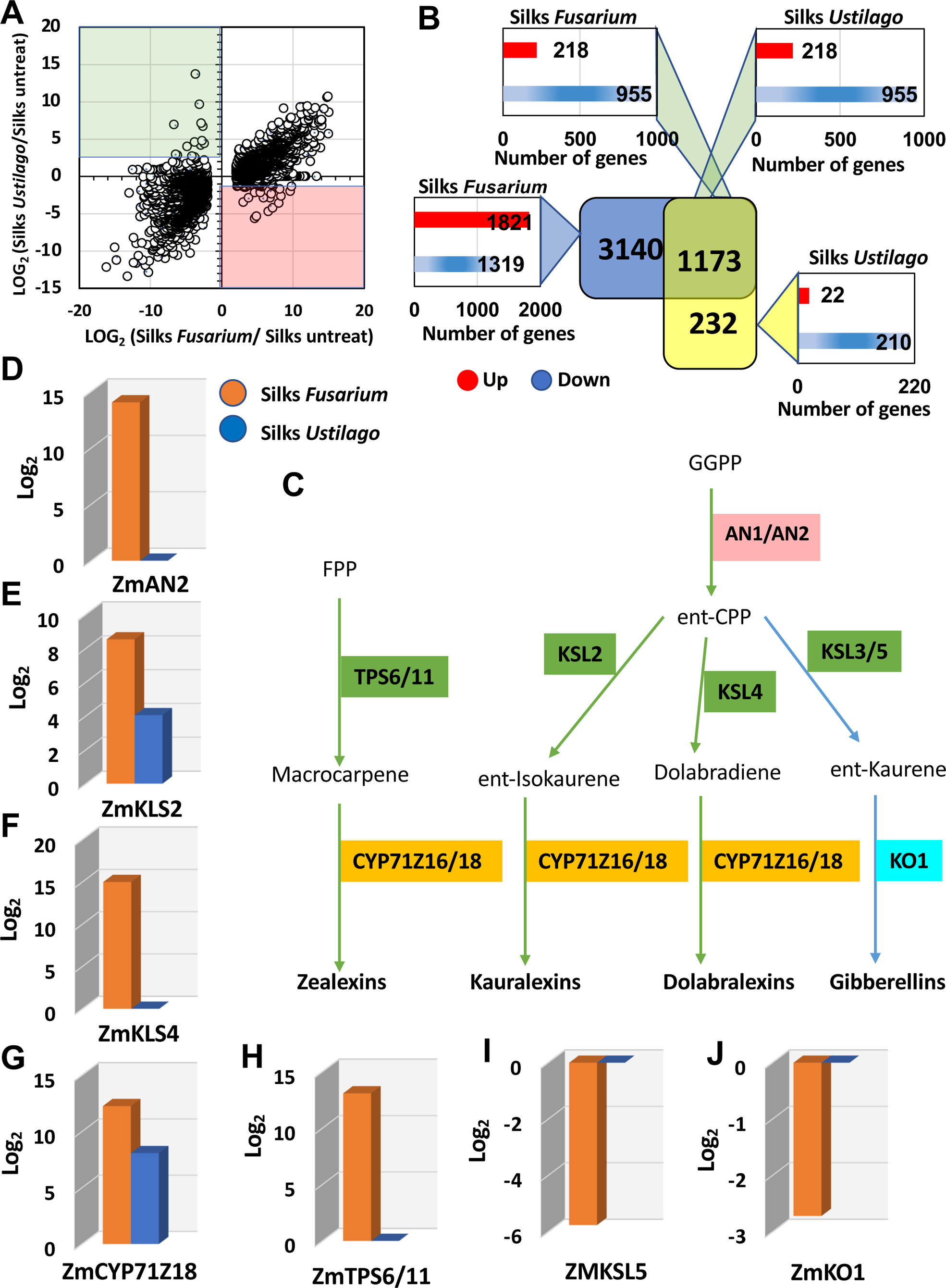
Activation of the defense system in maize silks to inhibit fungal growth. **(A)** Scatter plot representing the log_2_ for all DEGs in the maize silks infected with *F. graminearum* (x-axis) *versus* infected with *U. maydis* (y-axis) and **(B)** comparison of corresponding differentially expressed genes (DEGs). **(C)** Overview of the biosynthetic pathway of diterpenoids (zealexins, kauralexins and dolabralexins), major components of the maize defense system, and the growth hormone gibberellic acid. Class I diterpene synthases (diTPSs) are shown in red, class II diTPSs in green, and P450 enzymes in orange and cyan, respectively. Fold change expression analysis of the various diterpenoid biosynthesis enzyme genes namely **(D)** ent-copalyl diphosphate synthase (AN2), **(E)** ent-kaurene synthase2 (KSL2), **(F)** ent-kaurene synthase4 (KSL4), **(G)** cytochrome P450 monooxygenase (CYP71Z18), **(H)** diterpene synthases (TPS6/11), **(I)** ent-kaurene synthase5 (KSL5) and **(J)** *ent*-kaurene oxidase (KO1).

## Conclusions

First comparative studies on fungal and pollen tube invasion have shown among others that partly the same members of receptor-like kinase families like CrRLK1Ls and LRR RLKs, respectively, their ligands like RALFs and other small cysteine-rich proteins (CRPs) as well as membrane channels and ROS production/removal are involved in recognition and reception/rejection of the various biotic invaders (Kessler et al. 2010; Bircheneder and Dresselhaus 2016; Mondragón-Palomino et al. 2017; Zhou and Dresselhaus 2019; Gao et al. 2022). Much less is known about rejection and defense of unwanted pollen tubes and fungi after successful penetration. We have expanded previous studies using maize as a grass model species and characterized the molecular signatures of maize stigmas (silks) to four different invaders including own pollen tubes. The major findings are summarized in Figure 10. Although invasion occurs mainly via stigmatic papillae hair cell structures, further growth of each invader significantly varies triggering a different molecular response: growth of own pollen tubes occurs through the transmitting tract of the maize silk triggering regulation of relatively few genes including those involved in senescence processes, while *T. dactyloides* pollen tubes penetrate the same route, but often don’t target the transmitting tract and arrest after growing 2-3 cm triggering many responses including modifications of the cell wall of silk cells and defense reactions. Similar to pollen tubes, hyphae of *F. graminearium* grow initially intercellularly, until they penetrate and kill the cells. The invaded silk elicits the strongest response which appears to be mainly directed to inhibit fungal growth and to defend itself from the invader. In contrast, *U. maydis* grow exclusively intracellular without killing host cells. Compared to *F. graminearium,* the silk response is much weaker and can be considered to our opinion as a “half-hearted” defense reaction.

**Figure 10.**
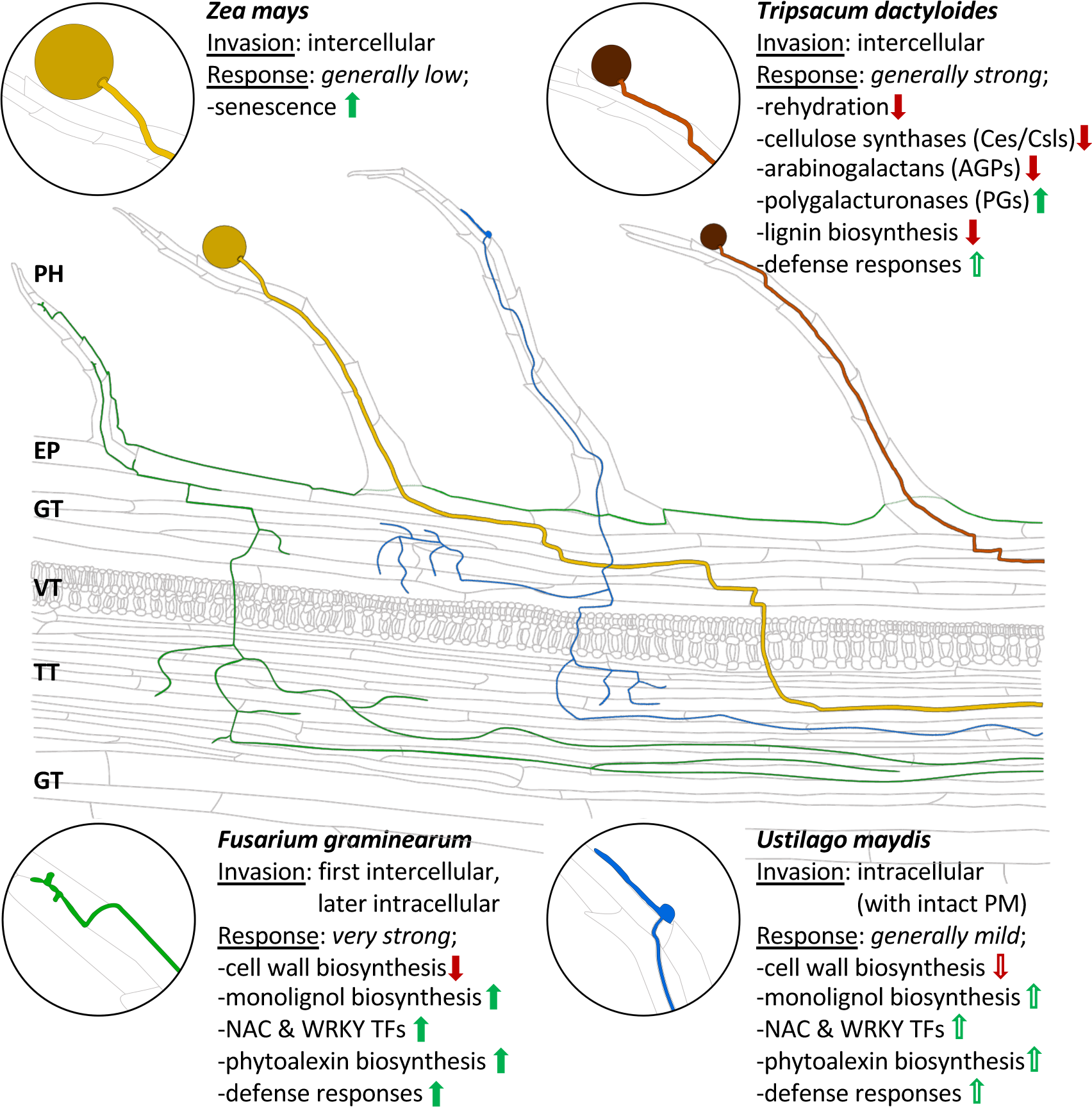
Summary of invasion pattern and major responses of maize silks to own and alien pollen tubes as well as to two different types of fungi. Ochre, own maize pollen tubes; brown, alien *T. dactyloides* pollen tubes; green, *F. graminearum and blue, U. maydis* hyphae. Enlargements show germinates pollen and fungal spores/conidia, respectively.

The molecular signatures associated to the various invaders now allows to target and enhance the specific molecular responses to defend invading fungal pathogens, but also to lower rejection processes during incompatible pollen tube invasion with the goal to overcome hybridization barriers and to increase the gene pool for plant breeding and seed production. The analysis and identification of the transcription factor regulatory network controlling cell wall development and defense responses in this study serves already as a very promising starting point for many future studies in grass crops to achieve plant pathogen resilience. Moreover, the comprehensive data set provided opens the windows also for other groups to identify, for example, more and novel genes involved in response to biotic invaders, to obtain stress-induced promoters and to identify further pathways involved in the described biotic interaction processes in maize and other grasses. Finally, we further propose that the highly elongated maize silk, which consists of only few cell types and is a relative transparent tissue, is also a very useful as a grass model for future pathogen studies.

## Material and methods

### Plant material and growth conditions

Seeds of maize (*Zea mays* L.) inbred line B73 as well as a *pUbi*::*CerTN-L15* marker line (see below) were germinated in germination traits containing soils and then transferred to pots (10 cm diameter, 10 seedlings per pot) containing a standard substrate and soil mixture (1:1, v/v) in the greenhouse (Begcy et al. 2019). After three weeks in the greenhouse, all genotypes were transferred to 10 L pots and grown under controlled conditions of 16 h of light at 26°C ± 2°C and 8 h of darkness at 22°C ± 2°C, and a constant air humidity of 60%. Supplemented light of ∼20,000 lux was provided to adjust day length duration. An automated temperature-water-based irrigation system was used to supply water according to plant consumption in a time-based preprogrammed schedule. Plants were fertilized twice a week with 2% (w/v) Hakaphos and monitored throughout their vegetative and reproductive development. Diploid *T. dactyloides* lines were grown in the greenhouse using the same growth conditions as maize. Plants were fertilized every second week.

Maize developmental stages were determined as previously described (Abendroth et al. 2011). Plant tissues were collected as follow: third leaves and whole roots were collected from maize seedlings at stage V3. Fresh pollen was collected at stage R1 (anthesis) (Begcy and Dresselhaus 2017) and either directly frozen or germinated for two hours on Petri dishes containing pollen germination medium (Begcy et al. 2019; Li et al. 2024). Silks at stage R1 were collected five days after emergence from ear husks. Silks of the same stage were harvested 8 hours after pollination (hap) using either pollen of maize inbred line B73 and diploid *T. dactyloides*, respectively. Whole developing seeds were excised 11 days after pollination (dap). After collection, all samples were immediately frozen in lipid N_2_ until further usage.

### Fungal material and growth conditions

*F. graminearum* wild-type strain 8/1 expressing GFP (Jansen et al. 2005) was used to visualize fungal growth in maize silks. Induction of conidia was carried out in wheat medium (7.5 g of 100% Bio-Weizengras LEBEPUR, Lebepur GmbH in 1 L of distilled water) on a shaker (150 rpm) in the dark at 28°C for 5 to 6 days and then filtered through a 70 μm sieve (Easytrainer, Greiner Bio-One). The filtrate was centrifuged at 4,000 rpm for 10 min at 4°C, and the supernatant was discarded. Inoculation medium for infection of maize cobs consisted of conidia resuspended in cold sterile distilled water to a final concentration of 1 to 2 x 10^5^ conidia per ml.

The GFP-expressing *U. maydis* strain used in this study was SG200-GFP, deriving from the solo-pathogenic wild-type strain SG200 (Kämper et al. 2006). *U. maydis* was grown on a shaker (200 rpm) at 28°C to an OD_600_ of 1 in YEPS-light medium (1% yeast extract, 0.4% peptone, and 0.4% sucrose), washed 3x with sterile distilled water and centrifuged at 3.500 rpm for 5 min each time (Woriedh et al. 2015). Spores were finally resuspended in sterile distilled water to an OD_600_ of 3 and the suspension was used for infecting plant material as described below.

### *In vitro* pollination of maize silks

Maize ears of approximately 8-10 cm in length from inbred line B73 were harvested and first cut longitudinally and then transversally (without wounding of silks) to obtain segments of about 5 cm in length. Segments were placed with the cut site on half MS-agar medium in small Petri dishes (5.5 cm diameter) that were then placed in bigger Petri dishes (13.5 cm diameter) lacking medium. All silks were orientated towards one direction, cut to the same length, and kept at room temperature before further processing. Silks were pollinated with freshly collected pollen from B73 as well as a homozygous transgenic line containing a *pUbi*::*CerTN-L15* marker construct (see below). Pollen tube growth was examined up to 24 hours after pollination (HAP) by confocal laser scanning microscopy.

### Fungal infection of maize silks

Maize silks were infected with the *F. graminearum* strain 8/1 and *U. maydis* strain SG200, both expressing GFP, using the silk channel inoculation method (Maier et al. 2006). In brief, fungal infection of maize silks was performed by applying 4 mL suspension with a syringe containing either *F. graminearum* conidia and *U. maydis* SG200-GFP spores, respectively. We monitored that silks were completely covered with the infection suspension solutions. After inoculation, samples were incubated at 28°C in the dark and cobs were covered with plastic bags for 3 days to favor fungal development. Infected silks were harvested 7 days after inoculation (DAI). A parallel set of untreated maize silks were harvested 7 days after silk emergence (DASE) as an ageing control. Aging and disintegration of silks is initiated about 7 DASE (Šimášková, et al. 2022). Similarly, maize silks inoculated with sterile water were used as mock infection control.

### Generation of a fluorescent maize pollen tube marker line

The calcium sensor line pDONR-CerTN-L15 (Denninger et al. 2014) containing cerulean and citrine fluorescent proteins was recombined into the p7i-Ubi-GW destination vector via LR reaction to generate *pUbi:*:*CerTN-L15*. This expression vector was transformed into maize HiIIA/B hybrid via Agrobacterium-mediated maize transformation as previously described (Frame et al. 2002). Transformants were selected by selection markers and fluorescent properties.

### Microscopic analyses

Confocal images were taken using a SP8 confocal laser scanning microscope (Leica). Cell walls were visualized using propidium iodide staining (Sigma-Aldrich). Infected maize silks were incubated with 50 mM propidium iodide staining solution for 15 min under dark conditions and mounted directly on objective slides before microscopy. Maize silks incubated with propidium iodide were excited at 488 nm and detected at 580-700 nm. GFP fluorescence was observed by exciting mounted silks samples with a 488 nm laser and emission detected at 500-540 nm (Antosz et al. 2020). Images were taken and processed using Leica Application Suite X (LAS X) software.

### RNA isolation and RT-qPCR

Total RNA was extracted using the Pure Link^TM^ RNA Mini Kit (Ambion, Life Technologies) and treated with RNase-Free DNase (Qiagen) according to the manufacturer’s instructions. Complementary DNA (cDNA) synthesis was performed using the Invitrogen SuperScript II reverse transcription system and oligo(dT) primers.

RNA quantification and quality control were assessed using a 2100 Bioanalyzer (Agilent Technologies). RT-qPCR reactions were carried out using KAPA SYBR Fast qPCR master mix (Peqlab) using an Eppendorf Mastercycler Realplex Real-Time detection system. Amplification efficiencies were calculated based on standard curves as previously described (Kim et al. 2021). Primer sequences with their corresponding melting temperatures and amplification efficiencies are provided in Supplementary Table S3. Cycling parameters consisted of 5 min at 95°C followed by 40 cycles of 95°C for 15 s, 60°C for 30 s, and 70°C for 30 s. Resulting Cq-values were imported into qbase+ v3.1 (Biogazelle). Quality control settings required that Cq-values from technical replicates differ by less than 0.5 cycles. Reactions were performed in triplicate for each RNA sample using at least three biological replicates. Normalization was based on three most stably expressed internal reference genes GRMZM2G000531, GRMZMG003734 and GRMZM2G003631 detected with the GeNorM program implemented by qBASE+ v3.1. The resulting calibrated normalized relative quantities (CNRQs) were then exported into Microsoft Excel for further analysis. Log_2_ transformed and calculation of fold change values was performed using the 2-ΔΔCt method.

### Library preparation and RNA-seq

Library preparation and RNA-seq were performed at the Regensburg University genomics service facility KFB (Competence Center for Fluorescent Bioanalytics) as described in the Illumina TruSeq Stranded mRNA Sample Preparation Guide, the Illumina HiSeq 1000 System User Guide (Illumina), and the KAPA Library Quantification Kit-Illumina/ABI Prism User Guide (Kapa Biosystems). Library construction was generated from cDNA fragments adenylated at their 3′-ends and ligated with indexing adapters, and specific cDNA libraries were subsequently created after PCR enrichment (Begcy et al. 2019). Libraries were quantified using the KAPA SYBR FAST ABI Prism Library Quantification Kit (Kapa Biosystems). Equimolar amounts of each library were used for cluster generation on the cBot using the Illumina TruSeq Paired-End Cluster Kit v3. Sequencing was performed on a HiSeq 1000 Illumina instrument using the indexed, 50 cycles paired-end read (PR) protocol and the TruSeq SBS v3 Reagents according to the Illumina HiSeq 1000 System User Guide. Image analysis and base calling resulted in bcl-files, which were converted into fastq-files using the bcl2fastq v2.18 software (https://www.illumina.com.cn/content/dam/illumina-support/documents/downloads/software/bcl2fastq/bcl2fastq2-v2-18-software-guide-15051736-01.pdf).

### Data processing, mapping, and differential expression

Paired-end reads were trimmed and filtered using Trimmomatic v.0.35 (Bolger et al. 2014). Data quality was assessed using FastQC. Processed paired-end reads were aligned to the high quality maize reference genome sequence AGPv3 assembly using annotation release-5b+ (Gramene AGPv3.27, https://www.maizegdb.org/assembly/) and a mapping program, STAR v. 2.5.2a (Dobin et al. 2013). Reads overlapping with more than one gene region were excluded from the analysis. To test for the presence/absence of batch effect and outliers, variance stabilizing transformation of raw counts and analyzed non-merged technical replicates was performed using R package DESeq2 (Love et al. 2014). For pairwise comparisons, DEGs were identified using DEseq2 by analyzing the number of reads aligned to the genes. The thresholds for differential expression were set at fold change >2 and P < 0.05 (after the false discovery rate adjustment for multiple testing) for the null hypothesis.

## Accession Numbers

A detailed list of genes and IDs mentioned in this study is provided in Supplemental Data Set 5.

## Availability of data and material

Data will be available in the NCBI Sequence Read Archive (SRA) under the bioproject accession number PRJNA1098880. All other materials are available from the corresponding authors upon request.

## Acknowledgments

We are grateful to Cathrin Kröger and Wilhelm Schäfer for providing the *F. graminearum* 8/1-GFP strain, to Vera Göhre and Michael Feldbrügge for the *U. maydis* SG200-GFP strain, and for their valuable support to optimize fungal growth conditions. We thank Mayada Woriedh for sample collection and Armin Hildebrand for plant care.

## Author contributions

TD designed the project. KB and TD conceived the experiments. KB, MMP and TD analyzed the data. KB and MMP performed the bioinformatics analyses. LZZ, PLS and MLM performed experimental assays. KB and TD wrote the article with input from all authors. All authors read and approved the article.

## Supplemental material

**Supplemental Table S1.** Summary of samples generated for this study, total reads per sample, and mapped reads aligned to the maize genome (AGPv3, Version 82.6).

**Supplemental Table S2.** List of differentially expressed genes (DEGs) in maize silks after self-pollination.

**Supplemental Table S3.** List of primers used in this study.

**Supplemental Figure S1.** Validation of expression pattern of 33 randomly selected genes from RNA-seq (right) data in eight of the 11 tissues investigated by using RT-qPCR (right). A Pearson correlation coefficient (R) >0.82 indicates a strong correlation.

**Supplemental Figure S2.** Hierarchical cluster analysis in response to pollination and fungal invasion. 18 clusters were identified by comparing the five silk tissues with various controls considering only expressed genes with TPM (transcript per million) values above 100.

**Supplemental Figure S3.** Key secondary wall-associated NAC transcription factor genes known as *ZmSWN*s are not up-regulated in maize silks in response to the various invaders investigated in this study. Expression levels are provided in transcript per million (TPM).

**Supplemental Figure S4.** MYB transcription factors gene expression in response to pollination and fungal invasion **(A)** Heat map displaying transcriptional expression of the entire MYB family. **(B)** Expression levels of ZmMYB20 **(C)** ZmMYB100 as examples for genes that are highly upregulated after Tripsacum pollination.

**Supplemental Figure S5.** Biological processes associated with fungal invasion in maize silks. **(A)** Biological processes associated with *F. graminearum* infection. **(B)** Biological processes associated with *U. maydis* infection.

**Supplemental Data Set 1.** Median expression values of treatment/tissue of genes mapped to the maize genome (AGPv3, Version 82.6).

**Supplemental Data Set 2.** Common DEGs between compatible and incompatible pollination.

**Supplemental Data Set 3.** Genes differentially expressed in response to incompatible pollination with *Tripsacum dactyloides*.

**Supplemental Data Set 4.** Downregulated rooted genes after incompatible pollination used for gene network analysis.

**Supplemental Data Set 5.** Detailed gene names and IDs described in this study.

## References

Abendroth LJ, Elmore, RW., Boyer, MJ., and Marlay, SK. Corn growth and development. 2011.

Abhinandan K, Sankaranarayanan S, Macgregor S, Goring DR, and Samuel MA. Cell–cell signaling during the Brassicaceae self-incompatibility response. Trends in Plant Science. 2022:27(5):472–487. 10.1016/j.tplants.2021.10.011

Amos RA and Mohnen D. Critical Review of Plant Cell Wall Matrix Polysaccharide Glycosyltransferase Activities Verified by Heterologous Protein Expression. Front Plant Sci. 2019:10:915. 10.3389/fpls.2019.00915

Antosz W, Deforges J, Begcy K, Bruckmann A, Poirier Y, Dresselhaus T, and Grasser KD. Critical Role of Transcript Cleavage in Arabidopsis RNA Polymerase II Transcriptional Elongation. Plant Cell. 2020:32(5):1449–1463. 10.1105/tpc.19.00891

Babu Y and Bayer M. Plant Polygalacturonases Involved in Cell Elongation and Separation—The Same but Different? Plants. 2014:3(4):613–623. 10.3390/plants3040613

Bassetti P and Westgate ME. Senescence and Receptivity of Maize Silks. Crop Science. 1993:33(2):275. 10.2135/cropsci1993.0011183X003300020012x

Begcy K and Dresselhaus T. Tracking maize pollen development by the Leaf Collar Method. Plant Reprod. 2017:30(4):171–178. 10.1007/s00497-017-0311-4

Begcy K, Nosenko T, Zhou L-Z, Fragner L, Weckwerth W, and Dresselhaus T. Male Sterility in Maize after Transient Heat Stress during the Tetrad Stage of Pollen Development. Plant Physiol. 2019:181(2):683–700. 10.1104/pp.19.00707

Bengoa Luoni S, Astigueta FH, Nicosia S, Moschen S, Fernandez P, and Heinz R. Transcription Factors Associated with Leaf Senescence in Crops. Plants. 2019:8(10):411. 10.3390/plants8100411

Bircheneder S and Dresselhaus T. Why cellular communication during plant reproduction is particularly mediated by CRP signalling. EXBOTJ. 2016:67(16):4849–4861. 10.1093/jxb/erw271

Boavida LC, Borges F, Becker JD, and Feijó JA. Whole Genome Analysis of Gene Expression Reveals Coordinated Activation of Signaling and Metabolic Pathways during Pollen-Pistil Interactions in Arabidopsis. Plant Physiology. 2011:155(4):2066–2080. 10.1104/pp.110.169813

Bolger AM, Lohse M, and Usadel B. Trimmomatic: a flexible trimmer for Illumina sequence data. Bioinformatics. 2014:30(15):2114–2120. 10.1093/bioinformatics/btu170

Bommert P and Whipple C. Grass inflorescence architecture and meristem determinacy. Seminars in Cell & Developmental Biology. 2018:79:37–47. 10.1016/j.semcdb.2017.10.004

Cao J, Liu H, Tan S, and Li Z. Transcription Factors-Regulated Leaf Senescence: Current Knowledge, Challenges and Approaches. IJMS. 2023:24(11):9245. 10.3390/ijms24119245

Cenci A, Chantret N, and Rouard M. Glycosyltransferase Family 61 in Liliopsida (Monocot): The Story of a Gene Family Expansion. Front Plant Sci. 2018:9:1843. 10.3389/fpls.2018.01843

Chase K, Belisle C, Ahlawat Y, Yu F, Sargent S, Sandoya G, Begcy K, and Liu T. Examining preharvest genetic and morphological factors contributing to lettuce (Lactuca sativa L.) shelf-life. Sci Rep. 2024:14(1):6618. 10.1038/s41598-024-55037-1

Chen L, Zhang L, Xiang S, Chen Y, Zhang H, and Yu D. The transcription factor WRKY75 positively regulates jasmonate-mediated plant defense to necrotrophic fungal pathogens. Journal of Experimental Botany. 2021:72(4):1473–1489. 10.1093/jxb/eraa529

Cheng X, Li M, Li D, Zhang J, Jin Q, Sheng L, Cai Y, and Lin Y. Characterization and analysis of *CCR* and *CAD* gene families at the whole-genome level for lignin synthesis of stone cells in pear (*Pyrus bretschneideri*) fruit. Biology Open. 2017:6(11):1602–1613. 10.1242/bio.026997

Denninger P, Bleckmann A, Lausser A, Vogler F, Ott T, Ehrhardt DW, Frommer WB, Sprunck S, Dresselhaus T, and Grossmann G. Male–female communication triggers calcium signatures during fertilization in Arabidopsis. Nat Commun. 2014:5(1):4645. 10.1038/ncomms5645

Ding Y, Murphy KM, Poretsky E, Mafu S, Yang B, Char SN, Christensen SA, Saldivar E, Wu M, Wang Q, et al. Multiple genes recruited from hormone pathways partition maize diterpenoid defences. Nat Plants. 2019:5(10):1043–1056. 10.1038/s41477-019-0509-6

Dobin A, Davis CA, Schlesinger F, Drenkow J, Zaleski C, Jha S, Batut P, Chaisson M, and Gingeras TR. STAR: ultrafast universal RNA-seq aligner. Bioinformatics. 2013:29(1):15–21. 10.1093/bioinformatics/bts635

Dong B, Yang Q, Song Z, Niu L, Cao H, Liu T, Du T, Yang W, Qi M, Chen T, et al. Hyperoside promotes pollen tube growth by regulating the depolymerization effect of actin-depolymerizing factor 1 on microfilaments in okra. Hortic Res. 2021:8(1):145. 10.1038/s41438-021-00578-z

Dorcey E, Urbez C, Blázquez MA, Carbonell J, and Perez-Amador MA. Fertilization-dependent auxin response in ovules triggers fruit development through the modulation of gibberellin metabolism in Arabidopsis. The Plant Journal. 2009:58(2):318–332. 10.1111/j.1365-313X.2008.03781.x

Dresselhaus T and Franklin-Tong N. Male–Female Crosstalk during Pollen Germination, Tube Growth and Guidance, and Double Fertilization. Molecular Plant. 2013:6(4):1018–1036. 10.1093/mp/sst061

Dresselhaus T, Lausser A, and Márton ML. Using maize as a model to study pollen tube growth and guidance, cross-incompatibility and sperm delivery in grasses. Annals of Botany. 2011:108(4):727–737. 10.1093/aob/mcr017

Entila F, Han X, Mine A, Schulze-Lefert P, and Tsuda K. Commensal lifestyle regulated by a negative feedback loop between Arabidopsis ROS and the bacterial T2SS. Nat Commun. 2024:15(1):456. 10.1038/s41467-024-44724-2

Frame BR, Shou H, Chikwamba RK, Zhang Z, Xiang C, Fonger TM, Pegg SEK, Li B, Nettleton DS, Pei D, et al. *Agrobacterium tumefaciens* -Mediated Transformation of Maize Embryos Using a Standard Binary Vector System. Plant Physiology. 2002:129(1):13–22. 10.1104/pp.000653

Gao Q, Wang C, Xi Y, Shao Q, Li L, and Luan S. A receptor–channel trio conducts Ca2+ signalling for pollen tube reception. Nature. 2022:607(7919):534–539. 10.1038/s41586-022-04923-7

Guo D-M, Ran J-H, and Wang X-Q. Evolution of the Cinnamyl/Sinapyl Alcohol Dehydrogenase (CAD/SAD) Gene Family: The Emergence of Real Lignin is Associated with the Origin of Bona Fide CAD. J Mol Evol. 2010:71(3):202–218. 10.1007/s00239-010-9378-3

He B, Shi P, Lv Y, Gao Z, and Chen G. Gene coexpression network analysis reveals the role of SRS genes in senescence leaf of maize (Zea mays L.). J Genet. 2020:99(1):3. 10.1007/s12041-019-1162-6

Huang H, Gehan MA, Huss SE, Alvarez S, Lizarraga C, Gruebbling EL, Gierer J, Naldrett MJ, Bindbeutel RK, Evans BS, et al. Cross-species complementation reveals conserved functions for EARLY FLOWERING 3 between monocots and dicots. Plant Direct. 2017:1(4). 10.1002/pld3.18

Jansen C, Von Wettstein D, Schäfer W, Kogel K-H, Felk A, and Maier FJ. Infection patterns in barley and wheat spikes inoculated with wild-type and trichodiene synthase gene disrupted *Fusarium graminearum*. Proc Natl Acad Sci USA. 2005:102(46):16892–16897. 10.1073/pnas.0508467102

Kämper J, Kahmann R, Bölker M, Ma L-J, Brefort T, Saville BJ, Banuett F, Kronstad JW, Gold SE, Müller O, et al. Insights from the genome of the biotrophic fungal plant pathogen Ustilago maydis. Nature. 2006:444(7115):97–101. 10.1038/nature05248

Kessler SA, Shimosato-Asano H, Keinath NF, Wuest SE, Ingram G, Panstruga R, and Grossniklaus U. Conserved Molecular Components for Pollen Tube Reception and Fungal Invasion. Science. 2010:330(6006):968–971. 10.1126/science.1195211

Kim T, Samraj S, Jiménez J, Gómez C, Liu T, and Begcy K. Genome-wide identification of heat shock factors and heat shock proteins in response to UV and high intensity light stress in lettuce. BMC Plant Biol. 2021:21(1):185. 10.1186/s12870-021-02959-x

Kodera C, Just J, Da Rocha M, Larrieu A, Riglet L, Legrand J, Rozier F, Gaude T, and Fobis-Loisy I. The molecular signatures of compatible and incompatible pollination in Arabidopsis. BMC Genomics. 2021:22(1):268. 10.1186/s12864-021-07503-7

Kumar A, Kanak KR, Arunachalam A, Dass RS, and Lakshmi PTV. Comparative transcriptome profiling and weighted gene co-expression network analysis to identify core genes in maize (Zea mays L.) silks infected by multiple fungi. Front Plant Sci. 2022:13:985396. 10.3389/fpls.2022.985396

Laitinen T, Morreel K, Delhomme N, Gauthier A, Schiffthaler B, Nickolov K, Brader G, Lim K-J, Teeri TH, Street NR, et al. A Key Role for Apoplastic H _2_ O _2_ in Norway Spruce Phenolic Metabolism. Plant Physiol. 2017:174(3):1449–1475. 10.1104/pp.17.00085

Lara-Mondragón CM and MacAlister CA. Arabinogalactan glycoprotein dynamics during the progamic phase in the tomato pistil. Plant Reprod. 2021:34(2):131–148. 10.1007/s00497-021-00408-1

Lausser A, Kliwer I, Srilunchang K, and Dresselhaus T. Sporophytic control of pollen tube growth and guidance in maize. Journal of Experimental Botany. 2010:61(3):673–682. 10.1093/jxb/erp330

Lauvergeat V, Lacomme C, Lacombe E, Lasserre E, Roby D, and Grima-Pettenati J. Two cinnamoyl-CoA reductase (CCR) genes from Arabidopsis thaliana are differentially expressed during development and in response to infection with pathogenic bacteria. Phytochemistry. 2001:57(7):1187–1195. 10.1016/S0031-9422(01)00053-X

Lee C, Teng Q, Zhong R, and Ye Z-H. Arabidopsis GUX Proteins Are Glucuronyltransferases Responsible for the Addition of Glucuronic Acid Side Chains onto Xylan. Plant and Cell Physiology. 2012:53(7):1204–1216. 10.1093/pcp/pcs064

Lee HK and Goring DR. Two subgroups of receptor-like kinases promote early compatible pollen responses in the *Arabidopsis thaliana* pistil. Journal of Experimental Botany. 2021:72(4):1198–1211. 10.1093/jxb/eraa496

Leydon AR, Beale KM, Woroniecka K, Castner E, Chen J, Horgan C, Palanivelu R, and Johnson MA. Three MYB Transcription Factors Control Pollen Tube Differentiation Required for Sperm Release. Current Biology. 2013:23(13):1209–1214. 10.1016/j.cub.2013.05.021

Li Q, Nie S, Li G, Du J, Ren R, Yang X, Liu B, Gao X, Liu T, Zhang Z, et al. Identification and Fine Mapping of the Recessive Gene BK-5, Which Affects Cell Wall Biosynthesis and Plant Brittleness in Maize. IJMS. 2022:23(2):814. 10.3390/ijms23020814

Li X, Bruckmann A, Dresselhaus T, and Begcy K. Heat stress at the bicellular stage inhibits sperm cell development and transport into pollen tubes. Plant Physiology. 2024:kiae087. 10.1093/plphys/kiae087

Li Y, Cheng X, Fu Y, Wu Q, Guo Y, Peng J, Zhang W, and He B. A genome-wide analysis of the cellulose synthase-like (Csl) gene family in maize. Biologia plant. 2019:63:721–732. 10.32615/bp.2019.081

Liang Y, Tan Z-M, Zhu L, Niu Q-K, Zhou J-J, Li M, Chen L-Q, Zhang X-Q, and Ye D. MYB97, MYB101 and MYB120 Function as Male Factors That Control Pollen Tube-Synergid Interaction in Arabidopsis thaliana Fertilization. PLoS Genet. 2013:9(11):e1003933. 10.1371/journal.pgen.1003933

Lorrai R and Ferrari S. Host Cell Wall Damage during Pathogen Infection: Mechanisms of Perception and Role in Plant-Pathogen Interactions. Plants. 2021:10(2):399. 10.3390/plants10020399

Love MI, Huber W, and Anders S. Moderated estimation of fold change and dispersion for RNA-seq data with DESeq2. Genome Biol. 2014:15(12):550. 10.1186/s13059-014-0550-8

Lu L, Hou Q, Wang L, Zhang T, Zhao W, Yan T, Zhao L, Li J, and Wan X. Genome-Wide Identification and Characterization of Polygalacturonase Gene Family in Maize (Zea mays L.). IJMS. 2021:22(19):10722. 10.3390/ijms221910722

Mafu S, Ding Y, Murphy KM, Yaacoobi O, Addison JB, Wang Q, Shen Z, Briggs SP, Bohlmann J, Castro-Falcon G, et al. Discovery, Biosynthesis and Stress-Related Accumulation of Dolabradiene-Derived Defenses in Maize. Plant Physiol. 2018:176(4):2677–2690. 10.1104/pp.17.01351

Maier FJ, Miedaner T, Hadeler B, Felk A, Salomon S, Lemmens M, Kassner H, and Schäfer W. Involvement of trichothecenes in fusarioses of wheat, barley and maize evaluated by gene disruption of the trichodiene synthase (Tri5) gene in three field isolates of different chemotype and virulence. Mol Plant Pathol. 2006:7(6):449–461. 10.1111/j.1364-3703.2006.00351.x

Mandrone M, Antognoni F, Aloisi I, Potente G, Poli F, Cai G, Faleri C, Parrotta L, and Del Duca S. Compatible and Incompatible Pollen-Styles Interaction in Pyrus communis L. Show Different Transglutaminase Features, Polyamine Pattern and Metabolomics Profiles. Front Plant Sci. 2019:10:741. 10.3389/fpls.2019.00741

Minic Z. Physiological roles of plant glycoside hydrolases. Planta. 2008:227(4):723–740. 10.1007/s00425-007-0668-y

Mitchell RAC, Dupree P, and Shewry PR. A Novel Bioinformatics Approach Identifies Candidate Genes for the Synthesis and Feruloylation of Arabinoxylan. Plant Physiology. 2007:144(1):43–53. 10.1104/pp.106.094995

Mól R, Filek M, Machackova I, and Matthys-Rochon E. Ethylene Synthesis and Auxin Augmentation in Pistil Tissues are Important for Egg Cell Differentiation after Pollination in Maize. Plant and Cell Physiology. 2004:45(10):1396–1405. 10.1093/pcp/pch167

Møller SR, Yi X, Velásquez SM, Gille S, Hansen PLM, Poulsen CP, Olsen CE, Rejzek M, Parsons H, Yang Z, et al. Identification and evolution of a plant cell wall specific glycoprotein glycosyl transferase, ExAD. Sci Rep. 2017:7(1):45341. 10.1038/srep45341

Mondragón-Palomino M, John-Arputharaj A, Pallmann M, and Dresselhaus T. Similarities between Reproductive and Immune Pistil Transcriptomes of *Arabidopsis* Species. Plant Physiol. 2017:174(3):1559–1575. 10.1104/pp.17.00390

Muhlemann JK, Younts TLB, and Muday GK. Flavonols control pollen tube growth and integrity by regulating ROS homeostasis during high-temperature stress. Proc Natl Acad Sci USA. 2018:115(47). 10.1073/pnas.1811492115

Murphy KM, Edwards J, Louie KB, Bowen BP, Sundaresan V, Northen TR, and Zerbe P. Bioactive diterpenoids impact the composition of the root-associated microbiome in maize (Zea mays). Sci Rep. 2021:11(1):333. 10.1038/s41598-020-79320-z

Nibbering P, Castilleux R, Wingsle G, and Niittylä T. CAGES are Golgi-localized GT31 enzymes involved in cellulose biosynthesis in Arabidopsis. The Plant Journal. 2022:110(5):1271–1285. 10.1111/tpj.15734

Ning Y and Wang G-L. Breeding plant broad-spectrum resistance without yield penalties. Proc Natl Acad Sci USA. 2018:115(12):2859–2861. 10.1073/pnas.1801235115

Ohtani M and Demura T. The quest for transcriptional hubs of lignin biosynthesis: beyond the NAC-MYB-gene regulatory network model. Current Opinion in Biotechnology. 2019:56:82–87. 10.1016/j.copbio.2018.10.002

Onelli E, Idilli AI, and Moscatelli A. Emerging roles for microtubules in angiosperm pollen tube growth highlight new research cues. Front Plant Sci. 2015:6. 10.3389/fpls.2015.00051

Park S-Y, Jauh G-Y, Mollet J-C, Eckard KJ, Nothnagel EA, Walling LL, and Lord EM. A Lipid Transfer–like Protein Is Necessary for Lily Pollen Tube Adhesion to an in Vitro Stylar Matrix. Plant Cell. 2000:12(1):151–163. 10.1105/tpc.12.1.151

Penning BW, Hunter CT, Tayengwa R, Eveland AL, Dugard CK, Olek AT, Vermerris W, Koch KE, McCarty DR, Davis MF, et al. Genetic Resources for Maize Cell Wall Biology. Plant Physiology. 2009:151(4):1703–1728. 10.1104/pp.109.136804

Pereira AM, Masiero S, Nobre MS, Costa ML, Solís M-T, Testillano PS, Sprunck S, and Coimbra S. Differential expression patterns of arabinogalactan proteins in Arabidopsis thaliana reproductive tissues. Journal of Experimental Botany. 2014:65(18):5459–5471. 10.1093/jxb/eru300

Qin L-X, Rao Y, Li L, Huang J-F, Xu W-L, and Li X-B. Cotton GalT1 Encoding a Putative Glycosyltransferase Is Involved in Regulation of Cell Wall Pectin Biosynthesis during Plant Development. PLoS ONE. 2013:8(3):e59115. 10.1371/journal.pone.0059115

Riglet L, Rozier F, Kodera C, Bovio S, Sechet J, Fobis-Loisy I, and Gaude T. KATANIN-dependent mechanical properties of the stigmatic cell wall mediate the pollen tube path in Arabidopsis. eLife. 2020:9:e57282. 10.7554/eLife.57282

Rozier F, Riglet L, Kodera C, Bayle V, Durand E, Schnabel J, Gaude T, and Fobis-Loisy I. Live-cell imaging of early events following pollen perception in self-incompatible Arabidopsis thaliana. Journal of Experimental Botany. 2020:71(9):2513–2526. 10.1093/jxb/eraa008

Sanchez AM, Bosch, M, Bots, M, Nieuwland, J, Feron, R, and Mariani, C. Pistil Factors Controlling Pollination. THE PLANT CELL ONLINE. 2004:16(suppl_1):S98–S106. 10.1105/tpc.017806

Schmelz EA, Kaplan F, Huffaker A, Dafoe NJ, Vaughan MM, Ni X, Rocca JR, Alborn HT, and Teal PE. Identity, regulation, and activity of inducible diterpenoid phytoalexins in maize. Proc Natl Acad Sci USA. 2011:108(13):5455–5460. 10.1073/pnas.1014714108

Sella Kapu NU and Cosgrove DJ. Changes in growth and cell wall extensibility of maize silks following pollination. Journal of Experimental Botany. 2010:61(14):4097–4107. 10.1093/jxb/erq225

Seo E, Choi D, and Choi. Functional studies of transcription factors involved in plant defenses in the genomics era. Briefings in Functional Genomics. 2015:14(4):260–267. 10.1093/bfgp/elv011

Shi D, Tang C, Wang R, Gu C, Wu X, Hu S, Jiao J, and Zhang S. Transcriptome and phytohormone analysis reveals a comprehensive phytohormone and pathogen defence response in pear self-/cross-pollination. Plant Cell Rep. 2017:36(11):1785–1799. 10.1007/s00299-017-2194-0

Showalter AM and Basu D. Glycosylation of arabinogalactan-proteins essential for development in Arabidopsis. Communicative & Integrative Biology. 2016:9(3):e1177687. 10.1080/19420889.2016.1177687

Šimášková, M, Daneva A, Doll N, Schilling N, Cubría-Radío M, Zhou L, De Winter F, Aesaert S, De Rycke R, Pauwels L, et al. KIL1 terminates fertility in maize by controlling silk senescence. The Plant Cell. 2022:34(8):2852–2870. 10.1093/plcell/koac151

Song Y, Jiang Y, Kuai B, and Li L. CIRCADIAN CLOCK-ASSOCIATED 1 Inhibits Leaf Senescence in Arabidopsis. Front Plant Sci. 2018:9:280. 10.3389/fpls.2018.00280

Stein O and Granot D. An Overview of Sucrose Synthases in Plants. Front Plant Sci. 2019:10:95. 10.3389/fpls.2019.00095

Szklarczyk D, Franceschini A, Wyder S, Forslund K, Heller D, Huerta-Cepas J, Simonovic M, Roth A, Santos A, Tsafou KP, et al. STRING v10: protein–protein interaction networks, integrated over the tree of life. Nucleic Acids Research. 2015:43(D1):D447–D452. 10.1093/nar/gku1003

Takenaka Y, Kato K, Ogawa-Ohnishi M, Tsuruhama K, Kajiura H, Yagyu K, Takeda A, Takeda Y, Kunieda T, Hara-Nishimura I, et al. Pectin RG-I rhamnosyltransferases represent a novel plant-specific glycosyltransferase family. Nature Plants. 2018:4(9):669–676. 10.1038/s41477-018-0217-7

Thompson M and Raizada M. Fungal Pathogens of Maize Gaining Free Passage Along the Silk Road. Pathogens. 2018:7(4):81. 10.3390/pathogens7040081

Vogel CM, Potthoff DB, Schäfer M, Barandun N, and Vorholt JA. Protective role of the Arabidopsis leaf microbiota against a bacterial pathogen. Nat Microbiol. 2021:6(12):1537–1548. 10.1038/s41564-021-00997-7

Watanabe Y, Schneider R, Barkwill S, Gonzales-Vigil E, Hill JL, Samuels AL, Persson S, and Mansfield SD. Cellulose synthase complexes display distinct dynamic behaviors during xylem transdifferentiation. Proc Natl Acad Sci USA. 2018:115(27). 10.1073/pnas.1802113115

Windari EA, Ando M, Mizoguchi Y, Shimada H, Ohira K, Kagaya Y, Higashiyama T, Takayama S, Watanabe M, and Suwabe K. Two aquaporins, SIP1;1 and PIP1;2, mediate water transport for pollen hydration in the *Arabidopsis* pistil. Plant Biotechnology. 2021:38(1):77–87. 10.5511/plantbiotechnology.20.1207a

Woriedh M, Merkl R, and Dresselhaus T. Maize EMBRYO SAC family peptides interact differentially with pollen tubes and fungal cells. EXBOTJ. 2015:66(17):5205–5216. 10.1093/jxb/erv268

Wu A, Hao P, Wei H, Sun H, Cheng S, Chen P, Ma Q, Gu L, Zhang M, Wang H, et al. Genome-Wide Identification and Characterization of Glycosyltransferase Family 47 in Cotton. Front Genet. 2019:10:824. 10.3389/fgene.2019.00824

Wu H, Xie D, Jia P, Tang Z, Shi D, Shui G, Wang G, and Yang W. Homeostasis of flavonoids and triterpenoids most likely modulates starch metabolism for pollen tube penetration in rice. Plant Biotechnology Journal. 2023:21(9):1757–1772. 10.1111/pbi.14073

Xu X, Yang Y, Liu C, Sun Y, Zhang T, Hou M, Huang S, and Yuan H. The evolutionary history of the sucrose synthase gene family in higher plants. BMC Plant Biol. 2019:19(1):566. 10.1186/s12870-019-2181-4

Xue J-S, Qiu S, Jia X-L, Shen S-Y, Shen C-W, Wang S, Xu P, Tong Q, Lou Y-X, Yang N-Y, et al. Stepwise changes in flavonoids in spores/pollen contributed to terrestrial adaptation of plants. Plant Physiology. 2023:193(1):627–642. 10.1093/plphys/kiad313

Yadav V, Wang Z, Wei C, Amo A, Ahmed B, Yang X, and Zhang X. Phenylpropanoid Pathway Engineering: An Emerging Approach towards Plant Defense. Pathogens. 2020:9(4):312. 10.3390/pathogens9040312

Yin J, Chang X, Kasuga T, Bui M, Reid MS, and Jiang C-Z. A basic helix-loop-helix transcription factor, PhFBH4, regulates flower senescence by modulating ethylene biosynthesis pathway in petunia. Hortic Res. 2015:2(1):15059. 10.1038/hortres.2015.59

Yin Y, Huang J, Gu X, Bar-Peled M, and Xu Y. Evolution of Plant Nucleotide-Sugar Interconversion Enzymes. PLoS ONE. 2011:6(11):e27995. 10.1371/journal.pone.0027995

Zabotina OA, Zhang N, and Weerts R. Polysaccharide Biosynthesis: Glycosyltransferases and Their Complexes. Front Plant Sci. 2021:12:625307. 10.3389/fpls.2021.625307

Zhang W, Zhao F, Jiang L, Chen C, Wu L, and Liu Z. Different Pathogen Defense Strategies in Arabidopsis: More than Pathogen Recognition. Cells. 2018:7(12):252. 10.3390/cells7120252

Zheng Y-Y, Lin X-J, Liang H-M, Wang F-F, and Chen L-Y. The Long Journey of Pollen Tube in the Pistil. IJMS. 2018:19(11):3529. 10.3390/ijms19113529

Zhong R, Lee C, McCarthy RL, Reeves CK, Jones EG, and Ye Z-H. Transcriptional Activation of Secondary Wall Biosynthesis by Rice and Maize NAC and MYB Transcription Factors. Plant and Cell Physiology. 2011:52(10):1856–1871. 10.1093/pcp/pcr123

Zhou L and Dresselhaus T. Friend or foe: Signaling mechanisms during double fertilization in flowering seed plants. In. Current Topics in Developmental Biology. (Elsevier), pp. 453–496. 10.1016/bs.ctdb.2018.11.013

Zhou L-Z, Juranić M, and Dresselhaus T. Germline Development and Fertilization Mechanisms in Maize. Molecular Plant. 2017:10(3):389–401. 10.1016/j.molp.2017.01.012

Zhou X, Liao H, Chern M, Yin J, Chen Y, Wang J, Zhu X, Chen Z, Yuan C, Zhao W, et al. Loss of function of a rice TPR-domain RNA-binding protein confers broad-spectrum disease resistance. Proc Natl Acad Sci USA. 2018:115(12):3174–3179. 10.1073/pnas.1705927115

Zhu J, Lolle S, Tang A, Guel B, Kvitko B, Cole B, and Coaker G. Single-cell profiling of Arabidopsis leaves to Pseudomonas syringae infection. Cell Reports. 2023:42(7):112676. 10.1016/j.celrep.2023.112676

